# A Dual Role for DGAT-mediated Lipid Droplet Biogenesis in Ferroptosis Regulation

**DOI:** 10.1101/2025.05.21.655263

**Authors:** Ana Kump, Leja Perne, Špela Koren, Eva Jarc Jovičić, Nastja Feguš, Carina Pinto Kozmus, Michele Wölk, Kristyna Brejchova, Maria Fedorova, Ondrej Kuda, Toni Petan

## Abstract

Lipid droplets (LDs) are dynamic fat storage organelles involved in fatty acid metabolism, signalling and trafficking. By storing polyunsaturated fatty acids (PUFAs) in the form of neutral lipids, LDs can either mitigate or exacerbate lipotoxic damage. However, the role of LDs in regulating cellular fatty acid distribution, membrane unsaturation and ferroptosis susceptibility remains poorly understood. Here, we show that inhibition of diacylglycerol acyltransferase (DGAT)-mediated LD biogenesis in PUFA-supplemented triple-negative breast cancer cells induces widespread lipidome rearrangements and membrane phospholipid acyl-chain remodelling, promoting lipid peroxidation and ferroptosis sensitivity. Lipidomic analyses reveal that LDs efficiently sequester exogenous PUFAs within triacylglycerols and cholesteryl esters, significantly altering the unsaturation profiles of these neutral lipids. When LD formation is impaired by DGAT inhibition, PUFAs are redistributed into membrane ester and ether glycerophospholipids, enhancing overall membrane unsaturation, lipid peroxidation, and increasing ferroptosis susceptibility, even in the absence of additional ferroptosis inducers. In contrast, in ferroptosis- and PUFA-resistant lung cancer cells, LDs exhibit a dual role, whereby the mode of ferroptosis induction and PUFA loading determined whether DGAT inhibition promoted or protected against ferroptosis. The pro-ferroptotic function of LDs predominates in these cells, particularly under conditions of ferroptosis suppressor protein 1 (FSP1) deficiency. This study highlights LDs as multifaceted regulators of ferroptosis sensitivity, integrating metabolic and redox quality control pathways.

## INTRODUCTION

Fatty acids (FAs) are essential for cellular structure and metabolism. Within cells, FAs are rarely found in their free form; instead, they are typically incorporated into complex lipids such as triacylglycerols (TGs), glycerophospholipids (PLs) and sterols. Their presence in membrane glycerophospholipids is crucial for the biophysical properties and dynamics of cellular membranes as well as the function of membrane proteins [1]. FAs also serve as energy sources, transcriptional regulators, signalling molecules and anchors for peripheral membrane proteins [2–5]. When cells fail to sufficiently metabolize or store excess FAs, they accumulate and cause lipotoxicity – disrupting cellular homeostasis, inducing oxidative stress, impairing organelle function and potentially triggering cell death [6]. FAs are primarily stored in their esterified forms as TGs and steryl esters within lipid droplets (LDs), dynamic organelles originating from the endoplasmic reticulum [7,8]. LDs undergo continuous cycles of biogenesis and breakdown, balancing FA storage and release in response to current cellular needs [9,10]. By sequestering excess FAs, LDs protect other organelles from lipotoxic damage, thus maintaining cellular homeostasis across various biological contexts [11–15]. Conversely, the regulated release of FAs from neutral lipids stored in LDs is crucial for fundamental processes such as energy production, membrane biogenesis and redox homeostasis [15,16]. However, the precise role of LDs in directing specific FAs to distinct cellular destinations for purposes like membrane lipid remodelling remains poorly understood.

The balance between saturated, unsaturated and polyunsaturated FAs (PUFAs) esterified in membrane PLs is crucial for proper membrane and organelle function [1]. PUFAs, such as docosahexaenoic acid (DHA; C22:6), are key membrane components necessary for the control of essential membrane properties, including bending, fluidity and thickness [17]. They are also substrates for controlled enzymatic oxygenation reactions, producing potent signalling molecules such as eicosanoids [18]. However, due to the weak bis-allylic methylene groups in their structure, PUFAs are also susceptible to peroxidation reactions, which can lead to membrane oxidative damage resulting in ferroptosis—a form of iron-dependent regulated cell death [19–22]. The oxidation of PUFA-containing lipids is essential for ferroptosis, which is triggered by the accumulation of membrane lipid peroxides and a reduced cellular capacity to eliminate them. Accordingly, lipid metabolic enzymes controlling the content of PUFAs in membrane phospholipids have been implicated in ferroptosis regulation [23–27]. Additionally, PUFA supplementation can result in ferroptosis in cultured cancer cells and dietary PUFA interventions compromise tumour growth by promoting ferroptosis [28–33].

The accumulation of PUFA-containing TGs within LDs can reduce lipotoxic damage, possibly by providing a protective environment within the dense neutral lipid LD core that prevents PUFA peroxidation [34,35]. This incorporation of PUFAs within LDs can also limit their availability for incorporation within membrane lipids, thereby reducing membrane lipid peroxidation and ferroptosis susceptibility [31,34–37]. On the contrary, the breakdown of PUFA-loaded LDs can also promote oxidative stress and ferroptotic cell death by delivering excess PUFAs to membrane phospholipids and other organelles [27,36,38–40]. Moreover, PUFA-rich neutral lipids within LDs can also be oxidized, promoting the initiation of ferroptosis [41]. Namely, in cells deficient in ferroptosis suppressor protein 1 (FSP1), PUFA-rich neutral lipids are readily oxidized and can be the source of lethal lipid peroxidation eventually leading to plasma membrane rupture and ferroptotic cell death [41]. Therefore, the role of LDs in modulating the cytotoxic and ferroptotic effects of PUFAs depends on multiple factors, including cell type-specific metabolic and redox programs, as well as environmental conditions that define lipid availability and cellular state.

We have previously shown that PUFA supplementation in triple-negative breast cancer cells exerts a dual effect, either promoting cell survival and proliferation or inducing oxidative stress and cell death, depending on concentration [36,42]. Here, we investigated the role of DGAT-mediated TG synthesis and LD biogenesis in lipidome remodelling, lipid peroxidation, and ferroptotic cell death in aggressive cancer cells supplemented with exogenous PUFAs. We found that suppressing LD biogenesis in PUFA-supplemented triple-negative breast cancer cells triggered widespread lipidome rearrangements and membrane acyl-chain remodelling, favouring lipid peroxidation and ferroptosis sensitivity in the absence of ferroptosis inducers. This ferroptosis-sensitive cellular state was characterized by enrichment of multiple PL classes with PUFAs. However, in lung cancer cells resistant to ferroptosis and PUFA-induced lipotoxicity, LDs played a context-dependent role, where the extent of PUFA-induced cell damage and mode of ferroptosis induction determined whether DGAT inhibition promoted or protected against ferroptosis.

## RESULTS

### DGAT inhibition potentiates PUFA-induced lipid peroxidation and ferroptosis

Our previous work demonstrated that PUFA supplementation induces LD accumulation in MDA-MB-231 triple-negative breast cancer cells, alongside effects ranging from protection against serum starvation to oxidative stress and cell death [36]. Here, we hypothesized that PUFA supplementation promotes membrane lipid peroxidation and ferroptosis, and that impairing LD biogenesis exacerbates these effects. To test this, we supplemented MDA-MB-231 cells with docosahexaenoic acid (DHA; C22:6, ω-3) and assessed the impact of DGAT inhibition on LD accumulation, oxidative stress, lipid reactive oxygen species (ROS) generation and cell viability.

DHA supplementation induced a concentration-dependent 2- to 3.5-fold increase in neutral lipid content (Fig. 1A), corresponding to substantial LD accumulation observed by confocal microscopy (Fig. 1B). To determine if DHA promotes LD biogenesis through DGAT enzymes, we used inhibitors targeting DGAT1 (T863) and DGAT2 (PF-06424439) (Fig. 1C). DGAT inhibition significantly reduced neutral lipid levels and fully abrogated DHA-induced LD accumulation (Figs. 1A, B). Next, we evaluated oxidative stress and lipid peroxidation using CM-H2DCFDA as a general ROS sensor and BODIPY 581/591 C11 as a lipid ROS sensor. DHA supplementation triggered a concentration-dependent rise in both lipid ROS and total ROS levels (Figs. 1D–F; Fig. S1A), effects that were further amplified by DGAT inhibition. Importantly, DGAT Inhibition potentiated DHA-induced cell death (Fig. 1G), indicating that LD biogenesis protects against PUFA-induced oxidative damage.

**Figure 1.**
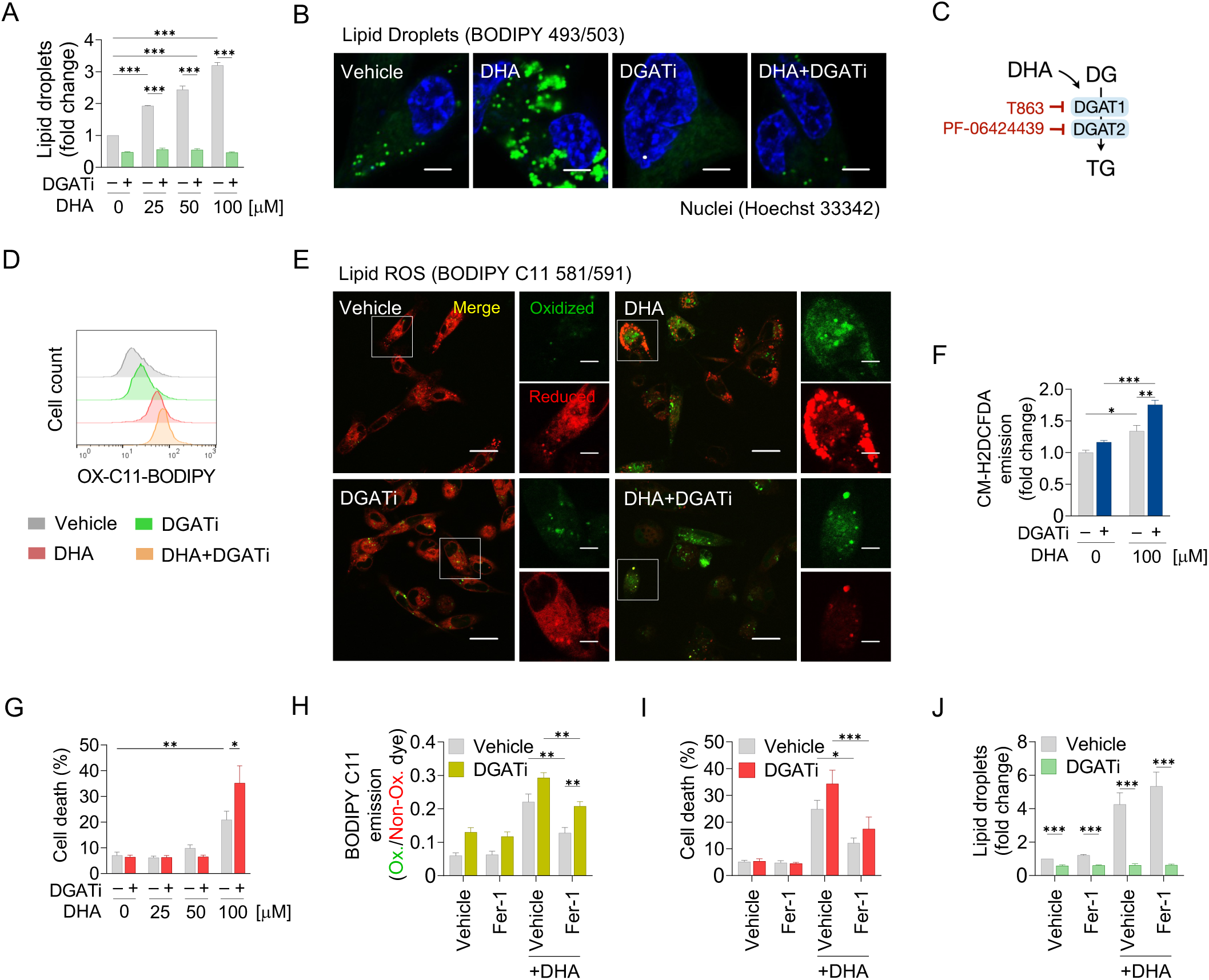
DGAT inhibition potentiates PUFA-induced lipid peroxidation and ferroptosis. (A) Lipid droplet levels measured by Nile Red staining in MDA-MB-231 cells treated with 25– 100 μM docosahexaenoic acid (DHA) for 24 h, with or without 20 μM DGAT1 and 20 μM DGAT2 inhibitors (DGATi). (B) Representative confocal microscopy images of MDA-MB-231 cells treated with 100 μM DHA for 24 h, with or without DGATi. Lipid droplets were stained by BODIPY 493/503 and nuclei by Hoechst 33342. Scale bar represents 5 μm. (C) General scheme of the last step in TG synthesis mediated by DGAT1 and DGAT2 enzymes, inhibited by specific inhibitors T863 and PF-06424439, respectively. (D) Lipid peroxidation assessed by BODIPY C11 581/591 probe using flow cytometry in MDA-MB-231 cells following the treatment described in (B). (E) Representative confocal microscopy images of MDA-MB-231 cells showing BODIPY C11 581/591 lipid peroxidation induced by the treatment described in (B). Scale bar represents 20 μm and 5 μm (inset). (F) Total reactive oxygen species (ROS) measured by CM-H2DCFDA in MDA-MB-231 cells following the treatment described in (B). (G) Cell death (%) determined by TMRM/YO-PRO-1 staining in MDA-MB-231 cells following the treatment described in (A). (H) Lipid peroxidation measured by BODIPY C11 581/591 in MDA-MB-231 cells treated with 100 μM DHA and/or 5 μM ferrostatin-1 (Fer-1) for 24 h, with or without DGATi. (I) Cell death (%) determined by TMRM/YO-PRO-1 staining in MDA-MB-231 cells following the treatment described in (H). (J) Lipid droplet levels measured by Nile Red staining in MDA-MB-231 cells following the treatment described in (H). All data are presented as mean ± SEM of at least three independent biological replications (significance calculated by two-way ANOVA with Tukey adjustment). p > 0.05 = ns; ^∗^p < 0.05; ^∗∗^p < 0.01; ^∗∗∗^p < 0.001.

To confirm that ferroptosis underlies DHA-induced cytotoxicity, we treated cells with ferrostatin-1, a radical-trapping ferroptosis inhibitor [43,44]. Ferrostatin-1 reduced DHA-induced lipid peroxidation (Fig. 1H) and cell death (Fig. 1I), without affecting LD levels (Fig. 1J), confirming ferroptosis as the dominant mode of death. Moreover, ferrostatin-1 attenuated the increase in lipid peroxidation (Fig. 1H) and cell death (Fig. 1I) caused by DGAT inhibition, further implicating LD biogenesis in modulation of ferroptosis susceptibility. As an additional control, monounsaturated oleic acid (OA; C18:1, ω-9), known to resist oxidation [45] promoted LD accumulation (Fig. S1B), but did not induce cell death, even with DGAT inhibition (Fig. S1C). Together, these results demonstrate that DGAT-mediated LD biogenesis buffers excess PUFAs and mitigates lipid peroxidation, thereby protecting MDA-MB-231 cells from ferroptotic death.

### DHA supplementation remodels LD-stored neutral lipids

Our data show that DGAT inhibition, and the resulting impairment of LD biogenesis, sensitizes MDA-MB-231 cells to DHA-induced ferroptosis. A likely explanation is that blocking TG synthesis prevents sequestration of exogenous DHA into neutral lipids, enhancing its incorporation into membrane PLs and promoting lipid peroxidation (Fig. 2A). To test this, we analysed lipidome remodelling under low, sublethal DHA exposure (25 µM) to minimize lipotoxicity [36] (Fig. 1G). Cells were treated with DHA for 1, 4 and 24 hours, with or without DGAT inhibitors, followed by LC-MS/MS-based lipidomics.

**Figure 2.**
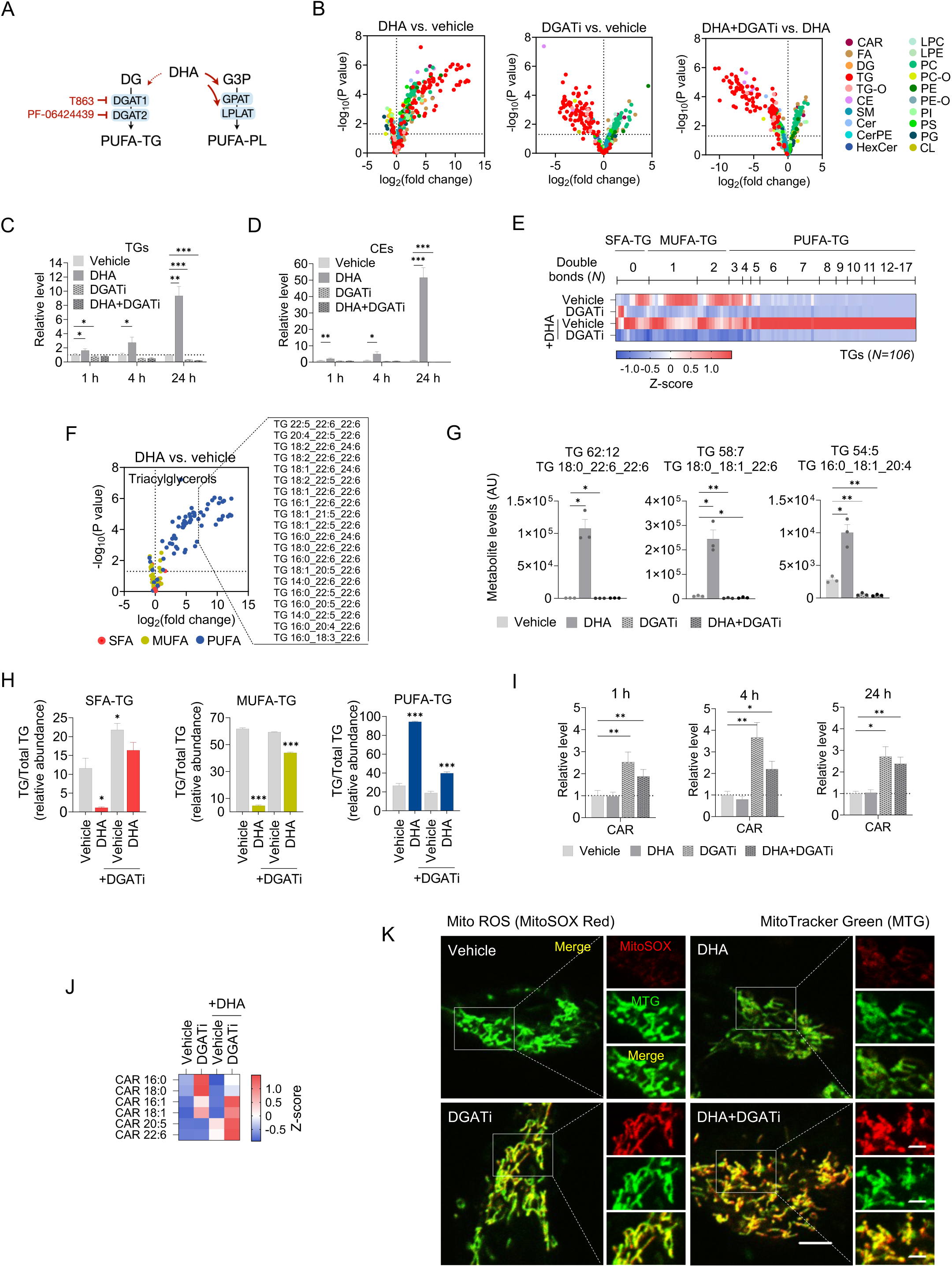
DHA supplementation remodels LD-stored neutral lipids. (A) Model illustrating the proposed mechanism of DGAT-mediated sensitization to PUFA-induced ferroptosis. (B) Volcano plots showing lipids altered by 24 h treatment of MDA-MB-231 cells with 25 μM DHA, 20 μM DGAT1 and 20 μM DGAT2 inhibitors (DGATi) and a combination of both treatments. Lipids are coloured by lipid class as indicated in the legend. Data were generated from three independent biological replicates using log_2_-transformed fold-change values and multiple unpaired *t*-tests (p < 0.05). (C, D) Relative total levels of TGs (C) and CEs (D) in MDA-MB-231 cells treated with 25 μM DHA for 1, 4 and 24 h, with or without DGATi. Data are presented as mean ± SEM (unpaired *t*-test). p > 0.05 = ns; ^∗^p < 0.05; ^∗∗^p < 0.01; ^∗∗∗^p < 0.001. (E) Heatmap showing relative abundance of TGs (n = 106) in MDA-MB-231 cells following the treatment described in (B), sorted by degree of unsaturation (number of double bonds) and categorized as SFA-TG (containing only saturated FAs), MUFA-TGs (containing saturated and monounsaturated FAs) or PUFA-TG (containing polyunsaturated FAs with at least 2 double bonds per chain). (F) Volcano plot of TG species altered by 24 h treatment with 25 μM DHA in MDA-MB-231 cells. TGs are categorized and coloured by unsaturation as indicated in the legend. The 20 most upregulated TGs are highlighted and labelled. Data were generated using log_2_-transformed fold-change values and multiple unpaired *t*-tests (p < 0.05). (G) Abundance of selected PUFA-TGs in MDA-MB-231 cells following the treatment described in (B). Each point represents a biological replicate. Data are presented as mean ± SEM (unpaired *t*-test). p > 0.05 = ns; ^∗^p < 0.05; ^∗∗^p < 0.01; ^∗∗∗^p < 0.001. (H) Relative abundance of SFA-, MUFA-, or PUFA-containing TGs in MDA-MB-231 cells following the treatment described in (B). Data are presented as mean ± SEM (unpaired *t*-test). p > 0.05 = ns; ^∗^p < 0.05; ^∗∗^p < 0.01; ^∗∗∗^p < 0.001. (I) Relative total levels of CAR in MDA-MB-231 cells following the treatment described in (C). Data are presented as mean ± SEM (unpaired *t*-test). p > 0.05 = ns; ^∗^p < 0.05; ^∗∗^p < 0.01; ^∗∗∗^p < 0.001. (J) Heatmap showing the relative abundance of CARs (n = 6) in MDA-MB-231 cells after the same 24 h treatment as indicated in (B). Data were Z-score normalized. (K) Representative confocal microscopy images of MDA-MB-231 cells co-stained with MitoSOX Red and MitoTracker Green following treatment with 50 μM DHA for 24 h, with or without DGATi. Scale bar represents 5 μm and 2 μm (inset). Abbreviations: G3P = glycerol-3-phosphate, GPAT = glycerol-3-phosphate acyltransferase, LPLAT = lysophospholipid acyltransferase, PAP = phosphatidic acid phosphatase, CAR = acylcarnitine, FA = fatty acid, DG = diacylglycerol, TG = triacylglycerol, TG-O = ether-linked triacylglycerol, CE = cholesterol ester, SM = sphingomyelin, Cer = ceramide, CerPE = ceramide phosphoethanolamine, HexCer = hexosylceramide, LPC = lyso-phosphatidylcholine, LPE = lyso-phosphatidylethanolamine, PC = phosphatidylcholine, PC-O = ether-linked phosphatidylcholine, PE = phosphatidylethanolamine, PE-O = ether-linked phosphatidylethanolamine, PI = phosphatidylinositol, PS = phosphatidylserine, PG = phosphatidylglycerol, CL = cardiolipin.

Global lipidomic analysis across 565 species revealed substantial remodelling under all conditions, with 167, 267 and 302 significantly altered lipids in response to DGAT inhibition, DHA supplementation, or combined treatment, respectively (Fig. 2B). In DHA-supplemented cells, neutral lipids (TGs, CEs) were elevated, which was fully suppressed by DGAT inhibition (Fig. 2B). Time-course analysis confirmed progressive accumulation of TGs and CEs with DHA supplementation, and this was entirely blocked by DGAT inhibition (Figs. 2C, D). Focusing on TG acyl-chain remodelling, DHA supplementation preferentially increased TGs with five or more double bonds (Fig. 2E), including TGs containing one, two, or three PUFA chains (Figs. 2F, G). This confirmed efficient DHA incorporation into TGs, though other non-DHA-containing PUFA-TGs (e.g., arachidonic acid (AA)-containing), as well as several non-PUFA TGs, were also elevated (Figs. 2F, G; Fig. S2A). DGAT inhibition reduced the abundance of nearly all TG species, abolishing the DHA-induced accumulation (Figs. 2E, G; Figs. S2A–C). Similarly, DHA-induced PUFA-CE enrichment was completely blocked by DGAT inhibition (Figs. S2D, E). To assess global changes in TG unsaturation, we grouped TGs by acyl-chain unsaturation. DHA supplementation shifted TG composition from 26% to 94% PUFA-TGs (Fig. 2H; Fig. S2F), accompanied by declines in SFA-TGs (10% to 4%) and MUFA-TGs (60% to 6%), highlighting the remarkable plasticity of LDs in storing excess PUFAs. DGAT inhibition almost completely prevented this shift, depleting neutral lipid stores and blocking DHA incorporation into TGs and CEs.

Beyond neutral lipids, DHA supplementation modestly elevated diacylglycerols, sphingomyelins and unesterified DHA (Figs. S2G–I). DGAT inhibition further increased free DHA and diacylglycerols, suggesting persistent DHA excess and ongoing lipid remodelling (Figs. S2G, I). Strikingly, only DGAT inhibition induced a strong accumulation of acylcarnitines (Figs. 2I, J), particularly PUFA-containing acylcarnitines in DHA-supplemented cells (Fig. 2J; Fig. S2J). This buildup suggests mitochondrial FA import overload and dysfunction [46,47]. Consistent with this, DGAT inhibition increased mitochondrial ROS, which was further exacerbated by DHA supplementation (Fig. 2K). Mitochondrial fragmentation, another hallmark of mitochondrial dysfunction recently implicated in ferroptosis [48,49], was also evident in DHA-supplemented, DGAT inhibitor-treated cells (Fig. 2K). Altogether, these results indicate that blocking TG synthesis redirects excess DHA towards alternative pathways, including diacylglycerol and acylcarnitine production, driving elevated glycerophospholipid remodelling, mitochondrial stress and increased susceptibility to ferroptosis.

### Inhibition of LD biogenesis leads to lipidome remodelling favouring PUFA enrichment in phospholipids

To investigate whether PUFA incorporation into membrane PLs drives lipid peroxidation and ferroptosis sensitivity upon DGAT inhibition, we analysed PL acyl-chain remodelling following DHA supplementation and DGAT inhibition. Grouping PLs by unsaturation and using volcano plots we found that both DHA treatment and DGAT inhibition increased many PUFA-PLs with various headgroups, each carrying one or two polyunsaturated acyl chains (Figs. 3A, B). Combining DHA supplementation with DGAT inhibition led to even greater PUFA-PL enrichment (Fig. 3C). Heatmap analysis after z-score normalization revealed distinct patterns: DGAT inhibition primarily upregulated PUFA-PLs with 2–5 double bonds, including both ester- (Fig. 3D) and ether-linked species (Fig. 3E). DHA supplementation alone increased PLs with 5–7 double bonds, while DHA combined with DGAT inhibition markedly increased a broader set of highly unsaturated PLs with 6–12 double bonds, particularly in ester-linked species (Figs. 3D, E). Tracking the relative abundances of PLs grouped by their unsaturation revealed that PUFA-PL levels increased over time with DHA supplementation, and DGAT inhibition further raised PUFA-PLs in both control and in DHA-supplemented cells (Figs. 3F, G; Fig. S3A). Interestingly, DHA supplementation also raised fully saturated SFA-PLs. Together, these results show that DGAT inhibition and DHA supplementation synergistically remodel the membrane lipidome toward PUFA enrichment.

**Figure 3.**
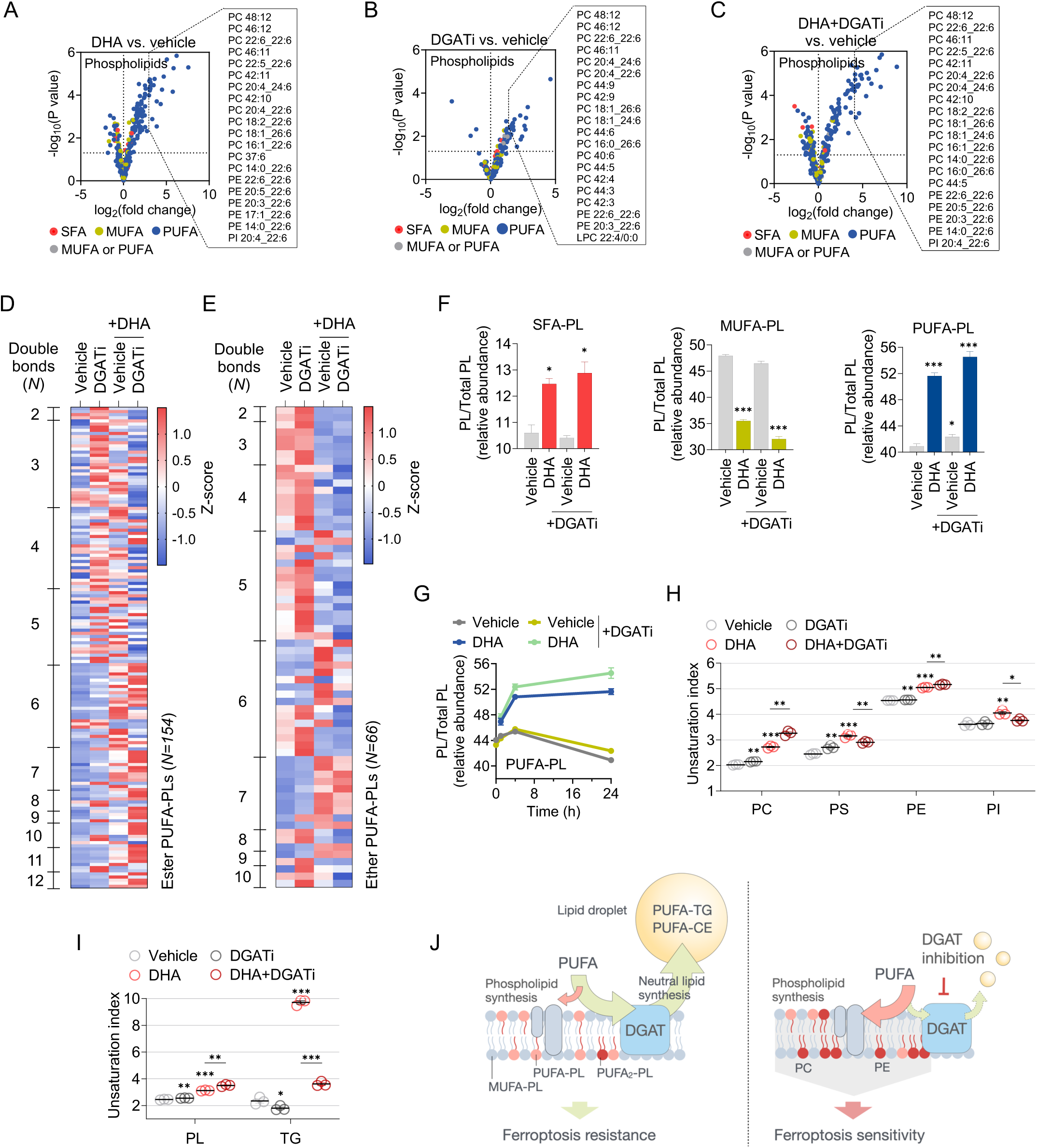
Inhibition of LD biogenesis leads to lipidome remodelling favouring PUFA enrichment in phospholipids. (A–C) Volcano plot of phospholipids (PLs) altered by 24 h treatment of MDA-MB-231 cells with 25 μM docosahexaenoic acid (DHA), with or without 20 μM DGAT1 and 20 μM DGAT2 inhibitors (DGATi). PLs were categorized and coloured as SFA (containing 0 double bonds in total), MUFA (containing 1 double bond in total), PUFA (containing > 2 double bonds in total), MUFA or PUFA (2 double bonds in total) as indicated in the legend. Data from three independent biological replications were prepared using log_2_-transformed fold-change values and multiple unpaired *t*-tests (p < 0.05). (D) Heatmap showing relative abundances of all quantified ester-linked PLs that contain polyunsaturated (≥ 2 double bonds) fatty acyl chains (PUFA-PLs) (n = 155) following the treatment described in (A). Lipids are sorted by degree of unsaturation (number of double bonds). Data were Z-score normalized. (E) Heatmap showing relative abundances of all quantified ether-linked PLs that contain polyunsaturated (≥ 2 double bonds) fatty acyl chains (PUFA-PLs) (n = 66) following the treatment described in (A). Lipids are sorted by degree of unsaturation (number of double bonds). Data were Z-score normalized. (F) Relative abundance of SFA-, MUFA-, and PUFA-containing PLs following the treatment described in (A). Data are presented as mean ± SEM (unpaired *t*-test). p > 0.05 = ns; ^∗^p < 0.05; ^∗∗^p < 0.01; ^∗∗∗^p < 0.001. (G) Relative abundance of PUFA-containing PLs following 1, 4 and 24 h treatment described in (A). Data are presented as mean ± SEM. (H) Abundance-weighed unsaturation index of the major PL classes following the treatment described in (A). Each point represents a biological replicate. Data are presented as mean ± SEM (unpaired *t*-test). p > 0.05 = ns; ^∗^p < 0.05; ^∗∗^p < 0.01; ^∗∗∗^p < 0.001. (I) Abundance-weighed unsaturation index of the PL and TG lipid classes following the treatment described in (A). Each point represents a biological replicate. Data are presented as mean ± SEM (unpaired *t*-test). p > 0.05 = ns; ^∗^p < 0.05; ^∗∗^p < 0.01; ^∗∗∗^p < 0.001. (J) Model summarizing the effects of DGAT inhibition on lipid remodelling underlying ferroptosis sensitization in PUFA-supplemented cells. Abbreviations: PC = phosphatidylcholine, PS = phosphatidylserine, PE = phosphatidylethanolamine, PI = phosphatidylinositol, PUFA_2_-PL = phospholipid with two PUFA chains.

While PUFA-PE species are key ferroptosis drivers [50–52], other PL classes also undergo peroxidation [24,32,53–58]. To assess PUFA enrichment across different headgroups, we measured changes in the overall unsaturation within each PL class [59]. This analysis showed that DHA supplementation increased PUFA content across all PL classes, with the strongest enrichment observed in PC and PS (Fig. 3H). DGAT inhibition alone increased PC, PS, and, to a lesser extent, PE unsaturation. Notably, in DHA-treated cells, DGAT inhibition further increased PC and PE unsaturation but blunted PS and PI remodelling, driving a net rise in overall PL unsaturation (Fig. 3I). Lipid over-representation analysis (LORA) [60] confirmed specific enrichment of long-chain, highly unsaturated PC and PE species in DHA-and DGATi-treated cells (Fig. S3B). At the individual lipid level, multiple PUFA-PLs from all four PL classes containing one (PUFA_1_-PL) or two (PUFA_2_-PL) PUFA chains, including PE 18:0_22:6, PE 20:4_22:6, PC 22:6_22:6 and PI 20:4_22:6, were increased by DHA and further augmented by DGAT inhibition (Figs. S3C–E). Together, these findings show that ferroptosis sensitivity arises from broad remodelling across multiple PL classes, involving various PUFA-PLs with one or two PUFA chains. Overall, our lipidomics reveal that DHA supplementation drives extensive PL remodelling and unsaturation, and that DGAT inhibition further augments PUFA incorporation across different headgroups, promoting a ferroptosis-prone lipidome (Fig. 3J).

### DGAT inhibition fails to promote DHA-induced cytotoxicity in ferroptosis-resistant cells

To determine if the sensitizing effect of DGAT inhibition observed in ferroptosis-sensitive MDA-MB-231 cells extends to ferroptosis-resistant cells, we selected four cancer cell lines based on prior studies of ferroptosis sensitivity [23,58]. The selected cell lines displayed distinct responses in LD accumulation following DHA supplementation. A549 cells showed the most pronounced LD accumulation, with up to a 5-fold increase in neutral lipid levels (Figs. 4A, B; Fig. S4A). There was negligible rise in neutral lipid levels and LD content in HeLa cells, and a ∼3-fold increase in MCF7 and T-47D cells (Fig. 4B; Figs. S4A, B). DGAT inhibition consistently reduced LD levels across all cell lines (Figs. 4A, B; Figs. S4A, B). While DHA supplementation had no effect on lipid ROS in MCF7 cells (Fig. 4D; Figs. S4D, F), it caused a weak increase in A549 (Figs. 4C, D), HeLa (Fig. 4D; Figs. S4C, D) and T-47D cells (Fig. 4D; Figs. S4D, E). However, none of these cell lines showed significant changes in total ROS or cell viability following DHA supplementation alone or in combination with DGAT inhibitors (Figs. 4E, F). The modest rise in lipid ROS levels upon DHA supplementation in A549, HeLa and T-47D cells, did not result in cell death, which suggests that robust ferroptosis surveillance mechanisms may neutralize lipid peroxidation in these cells.

**Figure 4.**
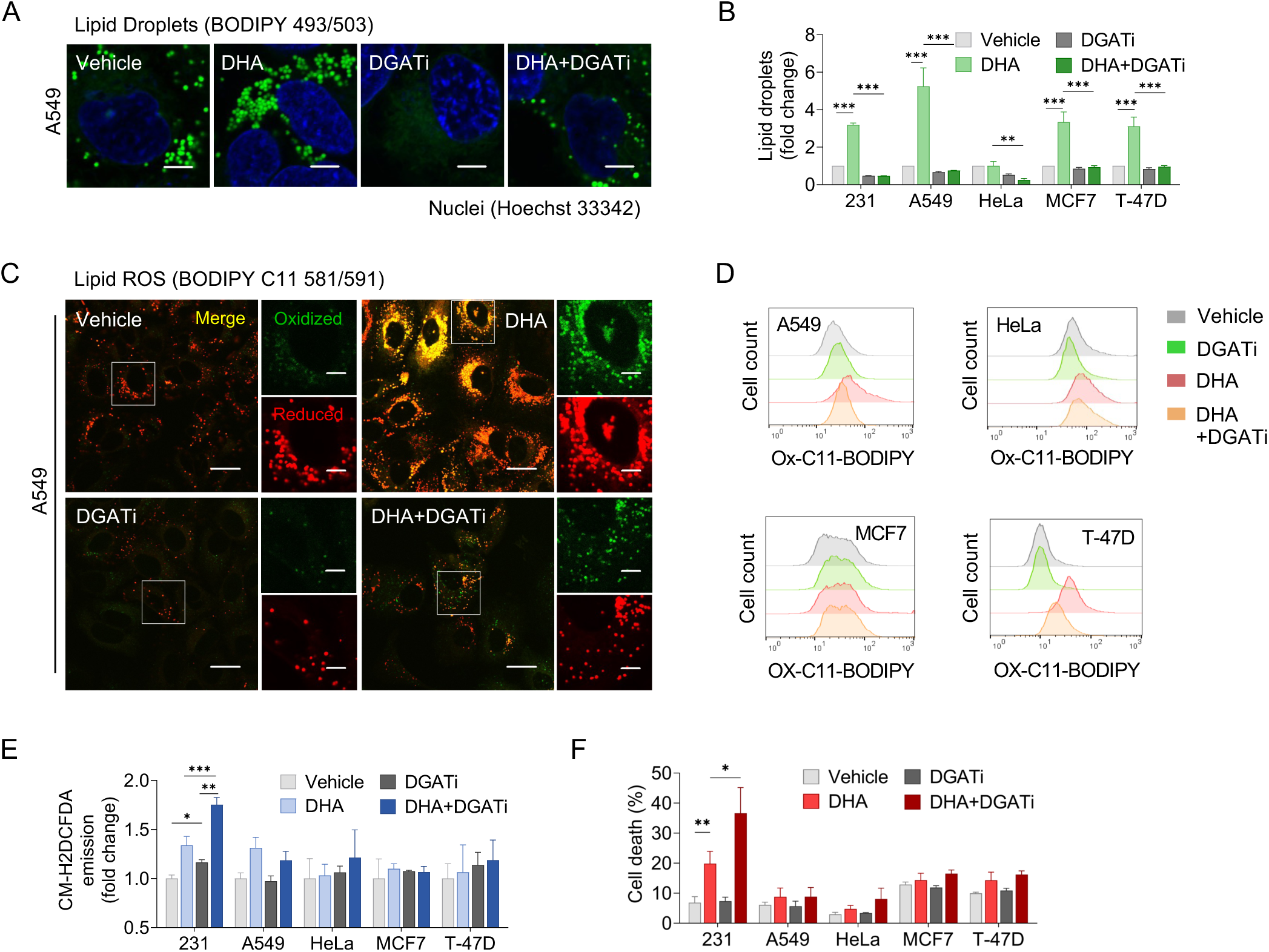
DGAT inhibition does not affect the viability of DHA-supplemented ferroptosis-resistant cells. (A) Representative confocal microscopy images of A549 cells treated with 100 μM docosahexaenoic acid (DHA) for 24 h, with or without 20 μM DGAT1 and 20 μM DGAT2 inhibitors (DGATi). Lipid droplets were stained by BODIPY 493/503 and nuclei by Hoechst 33342. Scale bar represents 5 μm. (B) Lipid droplet levels measured by Nile Red in indicated cell lines following the treatment described in (A). (C) Representative confocal microscopy images of A549 cells showing BODIPY C11 581/591 lipid peroxidation induced by the treatment described in (A). Scale bar represents 20 μm and 5 μm (inset). (D) Lipid peroxidation assessed by BODIPY C11 581/591 in indicated cell lines induced by the treatment described in (A). (E) Total reactive oxygen species (ROS) measured by CM-H2DCFDA in indicated cell lines following the treatment described in (A). (F) Cell death (%) determined by TMRM/YO-PRO-1 staining in indicated cell lines following the treatment described in (A). All data are presented as mean ± SEM of at least three independent biological replications (two-way ANOVA with Tukey adjustment). p > 0.05 = ns; ^∗^p < 0.05; ^∗∗^p < 0.01; ^∗∗∗^p < 0.001.

### Context-dependent effects of DGAT inhibition on ferroptosis sensitivity

Our findings in ferroptosis-resistant cell lines suggest that robust ferroptosis defence mechanisms prevent lethal lipid ROS buildup and may mask the role of LDs in modulating PUFA lipotoxicity. To unmask this potential role in A549 cells, we employed several strategies: altering DHA concentration, extending exposure duration, and impairing ferroptosis defence mechanisms (Fig. 5A). We first asked whether increasing the PUFA load alone could trigger cell death in A549 cells. Raising DHA levels to 200 µM induced significant cell death, even in the absence of ferroptosis inducers (Fig. 5B). Under these conditions, DGAT inhibition enhanced DHA-induced cell death. Mitochondrial ROS and fragmentation were strongly elevated in DHA-supplemented, DGAT-inhibited A549 cells (Fig. 5C), suggesting that mitochondrial dysfunction could contribute to cell death, consistent with observations in MDA-MB-231 cells (Fig. 2K). When this high PUFA load was combined with RSL3 treatment and FSP1 knockdown, we observed a synergistic induction of ferroptotic death (Fig. 5D; Figs. S5A, B), with earlier onset and greater intensity, which was further amplified by DGAT inhibition (Fig. 5E; Fig. S5B). Ferrostatin-1 rescued cells under these conditions, confirming ferroptotic cell death as the dominant mechanism (Fig. 5F). Together, these findings demonstrate that DGAT inhibition promotes ferroptosis in A549 cells under conditions of high PUFA overload.

**Figure 5.**
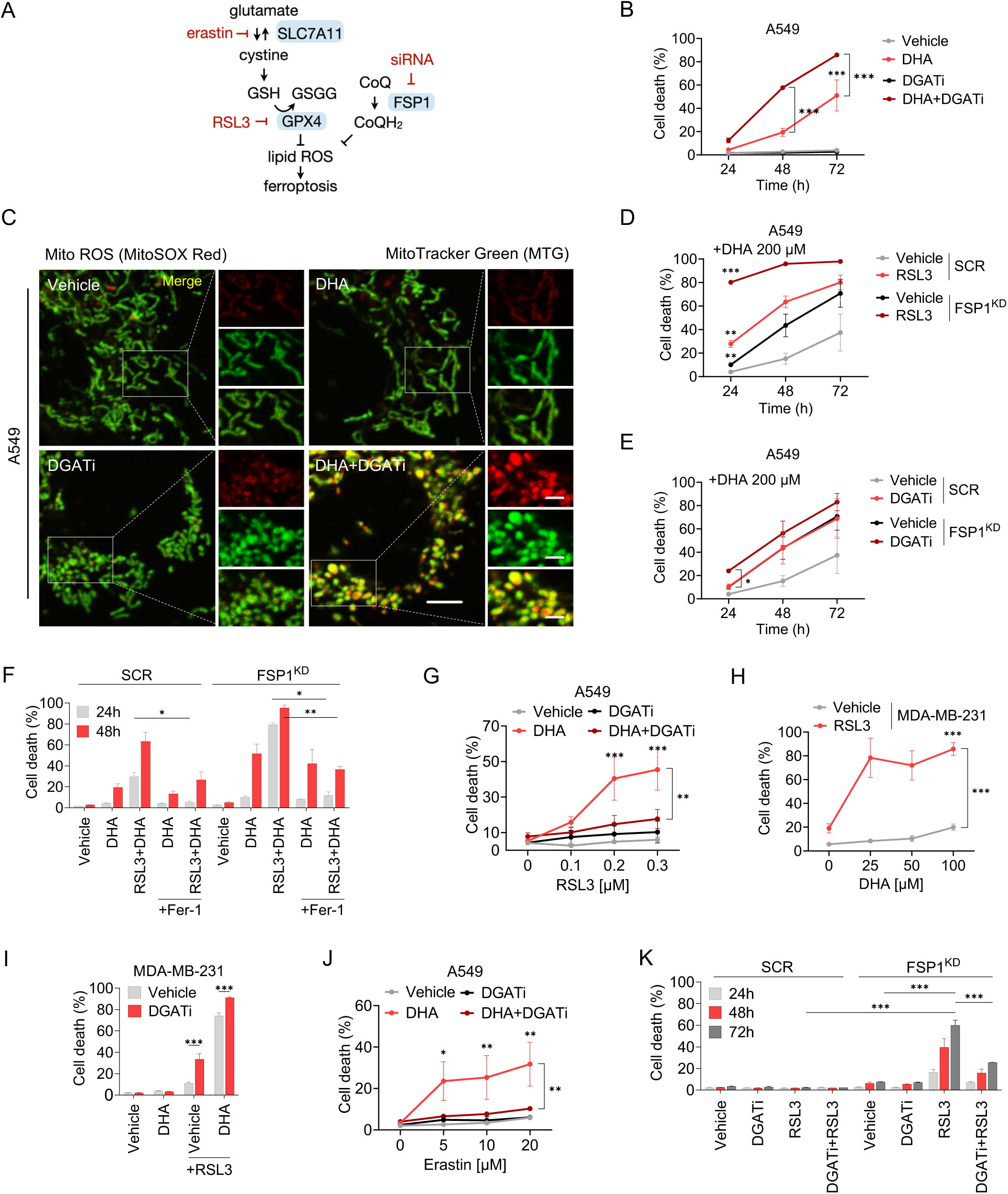
Context-dependent roles of LD biogenesis in ferroptosis. (A) Overview of key ferroptosis surveillance mechanisms and inhibitors/siRNAs used for induction of ferroptosis. (B) Cell death (%) in A549 cells treated with 200 μM docosahexaenoic acid (DHA) for 24, 48 and 72 h, with or without 20 μM DGAT1 and 20 μM DGAT2 inhibitors (DGATi). (C) Representative confocal microscopy images of A549 cells co-stained with MitoSOX Red and MitoTracker Green following treatment with 200 μM DHA for 24 h, with or without DGATi. Scale bar represents 5 μm and 2 μm (inset). (D, E) Cell death (%) in control (SCR) and FSP1-silenced (FSP1^KD^) A549 cells treated with 200 μM DHA for 24, 48 and 72 h, with or without: (D) 0.1 μM RSL3, and (E) DGATi. (F) Cell death (%) in SCR and FSP1^KD^ A549 cells treated with 200 μM DHA for 24 and 48 h, with or without 0.1 μM RSL3 and 5 μM ferrostatin-1 (Fer-1). (G) Cell death (%) in A549 cells treated with 100 μM DHA for 24 h, with or without DGATi, and 0.1, 0.2 or 0.3 μM RSL3. (H, I) Cell death (%) in MDA-MB-231 cells treated for 24 h with (H) 25, 50 or 100 μM DHA, with or without 0.05 μM RSL3, and (I) with 25 μM DHA, with or without DGATi and 0.05 μM RSL3. (J) Cell death (%) in A549 cells treated with 100 μM DHA for 24 h, with or without DGATi, and 5, 10 or 20 μM erastin. (K) Cell death (%) in SCR and FSP1^KD^ A549 cells treated for 24, 48 and 72 h with or without DGATi and 0.1 μM RSL3. Cell death was determined by TMRM/YO-PRO-1 staining and flow cytometry. All data are presented as mean ± SEM of at least two (B, F) or three independent biological replications. Significance was calculated by two-way ANOVA with Tukey adjustment (B, G–K) or unpaired *t*-test (D–F). p > 0.05 = ns; ^∗^p < 0.05; ^∗∗^p < 0.01; ^∗∗∗^p < 0.001. Abbreviations: SLC7A11 = solute carrier family 7 member 11, GSH = reduced glutathione, GSSG = oxidized glutathione, GPX4 = glutathione peroxidase 4, FSP1 = ferroptosis suppressor protein 1 (also known as AIFM2), CoQ = coenzyme Q (ubiquinone), CoQH_2_ = reduced coenzyme Q (ubiquinol), RSL3 = Ras Selective Lethal 3.

We next asked whether the role of LDs could also be revealed under lower PUFA load. Treatment of A549 cells with the GPX4 inhibitor RSL3 sensitized them to a non-toxic dose of DHA (100 µM), elevating both lipid ROS and cell death (Fig. 5G; Fig. S5C). Notably, in this context, DGAT inhibition reduced RSL3-induced lipid peroxidation and cell death (Fig. 5G; Figs. S5C, D), reversing its earlier pro-ferroptotic effect seen at higher PUFA levels and contrasting its role in the ferroptosis sensitive MDA-MB-231 cells. Indeed, even 50 nM RSL3 was sufficient to sensitize MDA-MB-231 cells to 25 µM DHA, and unlike in A549 cells, DGAT inhibition amplified this sensitisation (Figs. 5H, I; Fig. S5E). Corroborating the protective effect of DGAT inhibition observed in A549 cells, DGAT inhibition also reduced erastin-induced ferroptosis (Fig. 5J). To determine whether exogenous PUFA is required for this protective effect, we omitted DHA and induced ferroptosis by combining RSL3 with FSP1 knockdown. Even under these PUFA-poor, pro-ferroptotic conditions, DGAT inhibition also protected A549 cells from ferroptosis (Fig. 5K).

These findings support a model in which LDs can either promote or protect against ferroptosis, depending on the cellular context and PUFA load. Specifically, LDs promote ferroptosis in A549 cells under low PUFA exposure but protect under high PUFA overload, while in MDA-MB-231 cells, LDs consistently protect against ferroptosis.

### Cell-type-specific lipid ROS localization reflects LD function in ferroptosis

Our results suggest that the roles of LDs in lipid peroxidation and ferroptosis in A549 cells switch depending on the cellular context. Lange et al. [41] recently demonstrated in U2OS cells that PUFA-loaded LDs harbour oxidized neutral lipids and initiate ferroptosis in FSP1-deficient cells, even in the absence of GPX4 inhibitors [41]. They showed that LD-localized FSP1 prevents neutral lipid oxidation, thereby blocking LD-mediated promotion of ferroptosis. Inhibition of LD biogenesis under such conditions protected from ferroptosis.

Consistent with this mechanism, we found that FSP1 depletion sensitized DHA-overloaded A549 cells to ferroptosis even without RSL3 (Figs. 5E, F), suggesting that LD oxidation may also drive ferroptosis in these cells. In accordance with this idea, confocal imaging analyses demonstrated that LDs in A549 cells substantially colocalize with the oxidized form of the BODIPY C11 lipid ROS probe, suggesting the presence of oxidized lipids within LDs even under basal conditions (Figs. 6A, B). FSP1 depletion further increased LD-localized lipid ROS in both control and DHA-supplemented A549 cells, supporting its role in protecting LD-stored lipids from oxidation, as previously reported [41]. This finding also validated the use of the BODIPY C11 581/591 probe for estimating the oxidative conditions in LDs in our cellular system. The effect of FSP1 depletion on LD-localized lipid ROS was replicated in MDA-MB-231 cells. While colocalization of the oxidized probe with LDs was minimal under both basal and DHA-supplemented conditions, it increased upon FSP1 depletion (Figs. 6C, D).

**Figure 6.**
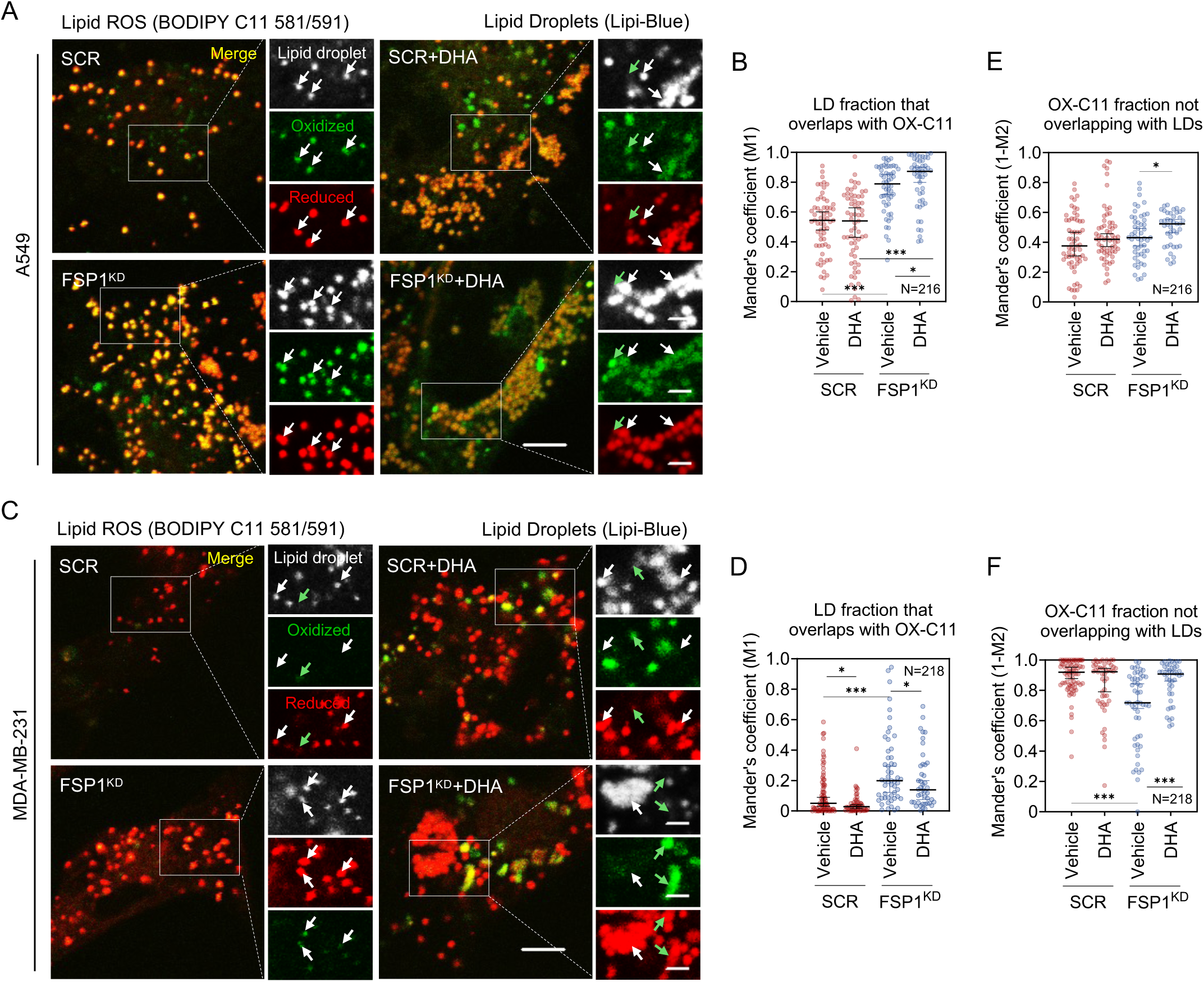
Lipid ROS localization reflects distinct roles of LDs in ferroptosis. (A) Representative confocal microscopy images of A549 cells co-stained with BODIPY C11 581/591 (lipid peroxidation sensor) and Lipi-Blue (lipid droplet marker), following FSP1 silencing (FSP1^KD^) and/or 200 μM docosahexaenoic acid (DHA) supplementation for 24 h. Colocalizations between lipid droplets and the oxidized dye are marked with white arrows. Green arrows mark oxidized probe signal that does not colocalize with lipid droplets. Scale bar represents 5 μm and 2 μm (inset). (B) M1 Mander’s coefficient quantifying colocalization between oxidized BODIPY C11 581/591 and Lipi-Blue in A549 cells as in (A). Each point represents an individual cell. (C) Representative confocal microscopy images of MDA-MB-231 cells co-stained with BODIPY C11 581/591 and Lipi-Blue, following FSP1^KD^ and/or 50 μM DHA supplementation for 24 h. Colocalizations between lipid droplets and the oxidized dye are marked with white arrows. Green arrows mark oxidized probe signal that does not colocalize with lipid droplets. Scale bar represents 5 μm and 2 μm (inset). (D) Same analysis as in (B) performed in MDA-MB-231 cells upon FSP1^KD^ and/or 50 μM DHA supplementation. (E) Quantification of oxidized BODIPY C11 581/591 outside lipid droplets in A549 cells as in (A). The complement of the M2 Mander’s coefficient (1–M2) was calculated to determine the proportion of lipid peroxidation signal not colocalizing with lipid droplets. Each point represents an individual cell. (F) Same analysis as in (E) performed in MDA-MB-231 cells upon FSP1^KD^ and/or 50 μM DHA supplementation. All data are presented as median ± 95% CI of two independent biological replications (unpaired *t*-test). p > 0.05 = ns; ^∗^p < 0.05; ^∗∗^p < 0.01; ^∗∗∗^p < 0.001.

Notably, DHA supplementation was not required for the FSP1 depletion-induced enhancement of LD-localized lipid ROS in either cell line, but DHA apparently increased the lipid ROS signal in non-LD cellular compartments (Figs. 6A, C). To quantify these shifts in lipid ROS localization, we calculated the complement of the M2 Manders coefficient, representing the fraction of oxidized lipid ROS signal outside LDs. This analysis confirmed that DHA-supplementation facilitated lipid ROS accumulation in non-LD compartments only in A549 cells, particularly upon FSP1 depletion (Figs. 6E, F). Together, these data indicate that LDs in A549 cells serve as oxidative hubs promoting the propagation of lipid peroxidation under both basal and PUFA-supplemented conditions. In contrast, LDs in MDA-MB-231 cells remain predominantly protective, maintaining a reduced environment, even under PUFA overload and FSP1 loss, suggesting they are not major sites of lipid oxidation in this context. In summary, our findings reveal significant cell-type differences in the regulation of lipid ROS accumulation within LDs. These differences suggest that the role of LDs in ferroptosis is influenced by LD-localized antioxidant mechanisms, including but not limited to FSP1.

## DISCUSSION

The capacity of LDs to sequester, store and release various FAs is central to their function as multifaceted organelles. By controlling FA availability and distribution, LDs are essential for supporting energy production or membrane synthesis as needed [9,15]. A less understood aspect of LD biology is their role in supplying specific FAs to maintain the diversity of membrane composition across different cellular contexts. In this study, we show that TG synthesis and LD biogenesis are critical components of the cellular response to PUFA supplementation, thereby preserving membrane and mitochondrial homeostasis, limiting lethal lipid ROS generation and preventing ferroptosis. Our data demonstrate that DGAT-mediated LD formation maintains lipidome balance by alleviating acyl-chain remodelling that underlies ferroptosis sensitivity. Inhibition of neutral lipid synthesis redirects excess PUFAs into membrane ether- and ester-linked PLs, resulting in elevated membrane unsaturation and increased abundance of pro-ferroptotic lipid species, such as PLs containing two polyunsaturated acyl chains [32,51,58]. This mechanism highlights a protective role for DGAT-driven LD biogenesis in mitigating PUFA-induced lipotoxicity and ferroptosis primarily by limiting PUFA incorporation into membranes. However, experiments in ferroptosis-resistant A549 lung adenocarcinoma cells reveal that this protective effect can be overridden. These cells accumulate lipid ROS in LDs even at basal conditions, and this increases with FSP1 depletion, supporting a mechanism of LD-driven ferroptosis, as seen in PUFA-supplemented U2OS cells [41]. In A549 cells, DHA sequestration into LDs appears to promote lipid ROS dispersion to other compartments, shifting the role of LDs from protective to pro-ferroptotic. Thus, our data underscore a context-dependent role of LD biogenesis in regulating PUFA lipotoxicity and ferroptosis sensitivity.

Our findings with triple-negative breast cancer cells suggest a protective role for DGAT-mediated TG synthesis and LD biogenesis against PUFA lipotoxicity and ferroptosis. This is based on our data showing that DGAT inhibition enhances lipid ROS generation and ferroptotic cell death induced by PUFA supplementation. Our lipidomic analyses further suggest that LD-driven lipidome remodelling underlies this protection. Neutral lipids stored in LDs substantially increase their unsaturation profiles upon PUFA supplementation, which points to a model of PUFA sequestration in TGs and CEs as a protective mechanism against PUFA-induced membrane and organelle damage in ferroptosis [34,35]. This model is supported by our data showing that impairment of TG synthesis by DGAT inhibition leads to enrichment of PLs with PUFAs. Specifically, we found that DGAT inhibition caused PUFA redistribution across different PLs classes, including ether- and ester-linked PC, PE, PS and PI species. These PLs contained either one or two polyunsaturated acyl chains, potentially contributing to ferroptosis susceptibility, as PLs from all of these four classes can be oxidized during ferroptosis [50–52,54–56]. Additionally, DGAT inhibition in DHA-supplemented cells caused a marked increase of PC and PE species containing two PUFA chains (PL-PUFA_2_). PC-PUFA_2_ species have been recently identified as drivers of ferroptosis, mediating the ferroptosis sensitizing effects of dietary PUFA-PCs or PUFAs [32,58]. On the other hand, PE lipids are significantly richer with PUFAs than PC species (Fig. 3H) and there is strong evidence for the involvement of both PE-PUFA_1_ and PE-PUFA_2_ species in ferroptosis [50–52,61,62]. The less abundant PS and PI species, which may also serve as substrates for peroxidation [53–56], were also enriched with PUFAs following DHA supplementation, with at least one PI-PUFA_2_ species containing AA and DHA elevated by DGAT inhibition. Nonetheless, the wide distribution of PUFA acyl chains across PL classes suggests that the general enrichment of membrane PLs could be the basis for ferroptosis sensitization in PUFA-treated MDA-MB-231 cells. This finding supports a model whereby the oxidation of PUFA acyl-chains in various phospholipids (or other complex lipids) collectively contribute to ferroptosis [63], but depending on their localization in the cell, the local redox balance and available repair mechanisms, they may or may not be involved in the initiation or progression of lipid peroxidation. In support of this model, the oxidation of PUFA acyl chains in neutral lipids stored in LDs can also initiate lipid peroxidation leading to ferroptosis [41]. The contribution of different lipid species and subcellular compartments to lipid peroxidation and ferroptosis across various contexts remains to be elucidated in future studies.

Our findings using the BODIPY-C11 lipid ROS probe, which serves as an estimate of the redox state of its environment [64], point to potential differences in LD oxidative state between MDA-MB-231 and A549 cells. In MDA-MB-231 cells, LDs primarily contained the reduced probe under basal conditions, and this was largely unaffected by DHA supplementation or FSP1 knockdown. In contrast, LDs in A549 cells accumulated the oxidized probe even under basal conditions, and following FSP1 knockdown, nearly all LDs in these cells became oxidized. This suggests the possibility that ferroptosis-sensitive MDA-MB-231 cells harbour LDs that are more resistant to oxidation, which is consistent with the model of PUFA sequestration in neutral lipids as a protective and predominant mechanism in these cells. Consequently, DGAT inhibition, which redirects PUFAs to membranes, becomes detrimental to these cells. Conversely, the predominantly oxidized LDs in the ferroptosis-resistant A549 cells may contribute to their resistance by limiting membrane lipid peroxidation even under basal conditions. However, when LD oxidation is further enhanced by FSP1 deficiency, it could promote the spread of lipid peroxidation, as observed in PUFA-supplemented U2OS cells [41]. Consistent with this LD-driven ferroptosis, our data suggests that DHA sequestration within LDs combined with FSP1 deficiency facilitates lipid ROS dispersion to other cellular compartments. This may explain why DGAT inhibition in A549 cells protects against RSL3- and FSP1-knockdown-induced ferroptosis and lower PUFA doses.

However, in A549 cells with compromised ferroptosis defences and exposed to very high PUFA loads, the role of LD biogenesis shifted from sensitization to protection from DHA-induced injury. This suggests that LDs serve multiple functions in ferroptosis, with their relative importance varying depending on cellular conditions. Under excessive PUFA load, the protective effect of PUFA sequestration in LDs, including prevention of membrane PUFA enrichment and mitochondrial dysfunction, may outweigh the pro-ferroptotic influence of LD oxidation. Additionally, very high PUFA levels likely induce various lipotoxic effects beyond ferroptosis, aligning with the broader role of LDs as primary FA sequestration sites that protect against lipotoxic stress. Therefore, our findings underscore a context-dependent role of LD biogenesis in the control of PUFA lipotoxicity and ferroptosis sensitivity depending on the metabolic and oxidative state of the cell.

Our results in DHA-supplemented MDA-MB-231 and A549 cells indicate that impaired TG synthesis redirects PUFAs towards mitochondria, leading to mitochondrial dysfunction. In MDA-MB-231 cells, the observed increase in polyunsaturated acylcarnitine levels suggests that this redirection ultimately exceeds mitochondrial capacity, thereby preventing efficient acylcarnitine import. These findings align with previous reports showing that DGAT inhibition causes mitochondrial damage due to acylcarnitine accumulation following FA overload triggered by starvation-induced autophagy [47]. Intact DGAT activity also supports mitochondrial homeostasis by regulating mitochondrial ROS levels and mitophagy, which is essential for clearing damaged mitochondria [65,66]. While PUFA supplementation alone caused only a modest increase in mitochondrial ROS in both cell lines, this effect was significantly amplified upon DGAT inhibition. The resulting ROS production likely contributes to ferroptosis, as shown in cells supplemented with PC-PUFA_2_ [32]. These lipids, which were upregulated by DHA supplementation and DGAT inhibition in our system, were recently shown promote mitochondrial ROS generation and ferroptosis initiation through interactions with ETC complex I [32]. Concurrently, we observed an increase in mitochondrial fragmentation, an indicator of mitochondrial dysfunction recently implicated in ferroptosis [48,49]. Specifically, the mitochondrial fission regulator dynamin-related protein 1 (Drp1) is activated during ferroptosis and promotes cell death execution through mitochondrial fragmentation [49]. Taken together, our data suggest that DGAT-mediated LD biogenesis supports mitochondrial homeostasis in DHA-supplemented cells, potentially preventing both ferroptosis and non-ferroptotic mitochondrial lipotoxicity [32,49,67,68].

Beyond ferroptosis, our data support the idea that cells tightly regulate membrane unsaturation to maintain a narrow range of physicochemical properties essential for membrane function and cellular fitness [59]. PUFA incorporation into membrane phospholipids increases membrane unsaturation, which can disrupt cellular homeostasis by altering key biophysical properties, such as membrane packing, bending and fluidity [17,69]. To counteract this, cells upregulate saturated PLs and cholesterol in membranes [59]. In our study, we observed a clear upregulation of saturated PLs and sphingolipids in DHA-supplemented cells, which likely helps maintain membrane rigidity following PUFA enrichment. This adaptive response may explain the need for a substantial plasticity of the LD lipidome, which—unlike cellular membranes—can significantly alter its unsaturation profile upon PUFA supplementation. It is thus possible that PUFA-induced membrane perturbations drive PUFA sequestration into LDs primarily to preserve membrane homeostasis, while also reducing the likelihood of membrane lipid peroxidation. Therefore, the sequestration of PUFAs into neutral lipids stored in LDs provides multiple benefits for the cell, including maintaining membrane function by regulating unsaturation levels and reducing ferroptosis susceptibility by limiting the availability of peroxidation-prone acyl chains in membranes.

This study identifies LDs as multifunctional hubs controlling ferroptosis through the management of cellular PUFA distribution, membrane diversity, lipid peroxidation, and mitochondrial integrity in human cancer cells. It remains unclear to what extent the identified mechanisms generalize to other cell types or in vivo contexts. While combined targeting of DGAT and ferroptosis surveillance mechanisms has therapeutic potential, the protective effect of DGAT inhibition in A549 cells suggests that promoting PUFA-rich LD accumulation may be more effective in some cancers. The dual roles of LDs within a single cell line underscore the need for caution when interpreting LD functions from limited conditions or screening data. The mechanisms underlying DGAT inhibition-induced PUFA redirection toward phospholipid remodelling remain to be defined. Moreover, this study addresses the role of LDs indirectly by studying the impact of their absence. Future work should explore how LD breakdown and interactions with other organelles contribute to ferroptosis and lipid peroxidation. It will be important to investigate how mitochondrial PUFA metabolism is affected by lipolysis, and how this influences ferroptosis. The coupling between LDs and peroxisomes in handling excess PUFAs also warrants further study. Finally, it will be exciting to uncover how LD breakdown pathways complement FSP1 quality control of LD lipids in ferroptosis.

## MATERIAL AND METHODS

### Chemicals

MDA-MB-231, T-47D, MCF7, A549, and HeLa human cancer cell lines were obtained from American Type Culture Collection (ATCC, USA). RPMI-1640 Medium (30-3001) and Eagle’s Minimum Essential Medium (EMEM) (30-2003) were obtained from ATCC (USA). Dulbecco’s Modified Eagle’s Medium/Nutrient Mixture F-12 (DMEM/F-12, 11330057), DMEM with high glucose and GlutaMAX supplement (DMEM/GlutaMAX, 61965-026), Dulbecco’s Phosphate-Buffered Saline (DPBS, 14190169), Fetal Bovine Serum (FBS, 10500064), Hanks’ Balanced Salt Solution (HBSS, 14025050), Hoechst 33342 (H3570), MitoTracker Green FM (M7514), MitoSOX Red (M36008), RPMI-1640 Medium without phenol red (11835063), TrypLE Select Enzyme (12563-029), L-Glutamine (25030081), Lipofectamine RNAiMAX Transfection Reagent (13778100) and Opti-MEM (11058021) were purchased from Thermo Fisher Scientific (USA). Human FSP1-targeting siRNAs and the AllStars Negative Control siRNA were from Qiagen (Germany). Docosahexaenoic acid (90310), oleic acid (90260), erastin (E7781), ferrostatin-1 (17729), (1S,3R)-RSL3 (19288) were obtained from Cayman Chemical (USA). BODIPY 493/503 (D3922), CM-H2DCFDA (C6827), BODIPY 581/591 C11 Lipid Peroxidation Sensor (D3861), Nile Red (N1142), Tetramethylrhodamine (TMRM) (T668), and YO-PRO-1 Iodide (Y3603) were from Thermo Fisher Scientific (USA). 7-aminoactinomycin D (7-AAD) (A9400), Insulin solution human (I9278), T863 (SML0539), PF-06424439 (PZ0233), essentially fatty acid-free (A7511)/fatty acid-free (A8806) bovine serum albumin (BSA) were purchased from Sigma-Aldrich (USA). Lipi-Blue (LD01) was purchased from Dojindo (Japan). DTT (R0862), Novex Tris-Glycine SDS 2X Sample Buffer (LC2676), EDTA-Free 100X Halt Protease Inhibitor Cocktail (78425), and Pierce 660 nm Protein Assay Kit (22662), were obtained from Thermo Fisher Scientific (USA). For Western blot (WB) analysis, WB-grade BSA from Sigma-Aldrich (USA) (A7030) was used. Nitrocellulose membrane (71224.01) was from SERVA (Germany), and Western Blocking Reagent (WBR) (11921673001) and Lumi-Light Western Blotting Substrate (12015196001) were from Roche Applied Science (Germany). β-actin primary antibodies (NB600-532) were from Novus Biologicals (UK), FSP1 primary antibodies (sc-37712) were from Santa Cruz Biotechnology (USA). Horseradish Peroxidase Conjugated Secondary Antibodies were from Jackson ImmunoResearch Laboratories (USA). All other chemicals were of at least analytical grade and purchased from Sigma-Aldrich (USA) or Serva (Germany).

### Cell Culture Conditions and Treatments

Cells were cultured in the following media, all supplemented with 10% FBS: RPMI-1640 was used for MDA-MB-231 cells; DMEM/F-12 for A549 cells; DMEM/GlutaMAX for HeLa cells; RPMI-1640 supplemented with 0.2 U/ml human insulin for T-47D cells; EMEM supplemented with 0.01 mg/ml human insulin for MCF-7 cells. All cells were maintained at 37 °C in a humidified atmosphere with 5% CO_2_, used at early passages, and passaged no more than 10 times. Mycoplasma contamination was routinely tested.

Fatty acids were stored in stock solutions (at concentrations higher than 10 mM) in absolute ethanol under argon at −80 °C. Prior to treatment, fatty acids were diluted in the appropriate culture medium with 10% FBS and incubated for 1 h at room temperature. DGAT1 and DGAT2 inhibitors (T863 and PF-07202654, respectively) were added to cells 2 h before treatments with fatty acids, ferrostatin-1, RSL3 or erastin and were present in the medium for the duration of the treatments. Appropriate vehicle controls (e.g., ethanol or DMSO) were included in all experiments to account for solvent effects.

### Gene Silencing Using Small-Interfering RNA

Reverse transfection was carried out in either 24-well, 6-well or 4-well plates using the following seeding densities: 3 × 10^4^ or 5 × 10^4^ A549 cells/well and 6 × 10^4^ MDA-MB-231 cells/well in 4- and 24-well plates; 2.5 × 10^5^ A549 cells/well and 3 × 10^5^ MDA-MB-231 cells/well in 6-well plates. Cells were transfected with a mixture of four FSP1-specific siRNAs (each at 5 nM, for a total concentration of 20 nM). A non-targeting control was included using 20 nM AllStars Negative Control siRNA. Transfection complexes were prepared using Lipofectamine RNAiMAX Transfection Reagent (1 µl/well for 4- and 24-well plates; and 7.5 µl/well for 6-well plates) and Opti-MEM medium, following the manufacturer’s instructions.

### Flow Cytometry Analysis

For all flow cytometry assays, cells were seeded into 24-well culture plates at a density of 3 × 10^4^ or 6 × 10^4^ cells per well and incubated for 24 h prior to treatment. Unless otherwise specified, cells were harvested after treatment, transferred to FACS tubes along with the culture supernatant, and centrifuged at 300 × *g* for 10 min. For neutral lipid analysis, cells were stained with 1 μg/ml Nile Red in DPBS for 10 min in the dark. Samples were analysed using excitation at 488 nm and an FL-1 emission filter (530/30 nm). At least 20,000 events were collected per sample using Cell Quest software (Becton Dickinson, USA). For cell death analysis, cells were stained with 25 nM TMRM in DPBS for 15 min in the dark, followed by staining with 50 nM YO-PRO-1 for an additional 10 min in the dark. Samples were diluted in DPBS containing 0.1% fatty acid-free BSA and analyzed by flow cytometry. Samples were analyzed using excitation at 488 nm, the FL-1 emission filter (530/30 nm) for YO-PRO-1 and the FL-3 filter (650LP) for TMRM. YO-PRO-1⁺/TMRM⁻ cells were gated as dead. At least 30,000 events were collected per sample. For lipid peroxidation analysis, cells were stained prior to treatment with 1 µM BODIPY 581/591 C11 in HBSS at 37 °C for 30 min. Following treatment, cells were harvested, centrifuged, resuspended in HBSS and analyzed by flow cytometry. Samples were analyzed with excitation set at 488 nm, an FL-1 emission filter (530/30 nm) for oxidized BODIPY C11 and FL-3 filter (650LP) for reduced BODIPY C11. At least 20,000 events were collected for each sample. For total ROS and cell death analysis, cells were stained with 1 µM CM-H2DCFDA in HBSS for 30 min, followed by a 2 h incubation in phenol red-free culture medium. Subsequently, cells were harvested, stained with 5 µg/ml 7-AAD for 10 min and analyzed by flow cytometry. Samples were analyzed using excitation at 488 nm, an FL-1 filter (530/30 nm) for CM-H2DCFDA and an FL-3 filter (650LP) for 7-AAD. ROS levels were determined in live cells (7-AAD^−^). At least 20,000 events were collected for each sample. For all flow cytometry assays, at least 2 independent biological replicates were performed.

### Live-Cell Imaging

Cells were seeded into 4-well glass bottom culture plates (Greiner Bio-One, 627870) at a density of 3 × 10^4^ or 6 × 10^4^ cells per well and incubated for 24 h. For lipid droplets and nuclei visualization, cells were washed with warm DPBS then stained for 15 min at 37 °C with 1 μM BODIPY 439/503 and 1 μg/ml Hoechst 33342 in culture medium without FBS, washed with warm DPBS and imaged. For visualization of mitochondria and mitochondrial ROS, cells were washed with warm HBSS then stained for 30 min at 37 °C with 100 nM MitoTracker Green and 1 μM MitoSOX Red in HBSS, washed with warm HBSS and imaged. For lipid peroxidation analysis, cells were washed with warm PBS and stained with 1 μM BODIPY 581/591 C11 in HBSS at 37 °C for 30 min prior to treatment. For lipid droplet and BODIPY C11 colocalization imaging, cells were stained with BODIPY 581/591 C11 prior to treatment as described above. Following treatment, cells were washed with warm DPBS, stained with 0.1 μM Lipi-Blue in culture medium without FBS for 15 min at 37 °C, washed with warm DPBS and imaged. Imaging was performed using a confocal laser scanning microscope (LSM 710; Carl Zeiss, Germany) with a stage-top CO_2_ incubation system (Tokai Hit, Japan). Images were processed and analyzed with ZEN (Carl Zeiss, Germany) and ImageJ (National Institute of Health, USA) software.

### Image Analysis

All image analysis was performed using Fiji [70] on at least 50 cells per sample. For colocalization quantification, confocal images were split into individual channels and processed side-by-side with the JaCoP plugin following the plugin instructions. Mander’s M1 and M2 coefficients were calculated individually for each cell using manual thresholding.

### Western Blot Analysis

Cells were seeded into 6-well culture plates at a density of 1.5 × 10^5^ or 3 × 10^5^ cells per well for 24 h. The medium was then replaced with fresh culture medium and cells were incubated for an additional 24 h. Cell lysates were prepared by washing cells with ice-cold DPBS and scraped into Tris-glycine SDS 2X sample buffer with 800 mM DTT and Halt Protease Inhibitor Cocktail. Lysates were incubated at 95 °C for 10 min and either stored on ice for immediate use or frozen at –80 °C for later analysis. Protein concentration was determined using the Pierce 660 nm Protein Assay Kit according to the manufacturer’s instruction. Equal amounts of protein (10 μg per sample) were loaded onto 10% SDS-PAGE gels and separated by electrophoresis. Proteins were then transferred onto nitrocellulose membranes using a wet western blot transfer system. Membranes were blocked for 1 h at room temperature in 1% Western Blocking Reagent (WBR) in TBS and incubated overnight at 4 °C with primary antibodies diluted in 0.5% WBR in 0.1% TBS-T (TBS with 0.1% Tween-20). After three washes with TBS-T, membranes were incubated with horseradish-peroxidase-labelled secondary antibodies diluted 1:10,000 in 0.5% WBR in TBS-T for 1 h at room temperature. Membranes were washed three times in TBS-T, signal detection was performed using Lumi-Light Western Blotting Substrate, and chemiluminescent signals were visualized using a Gel Doc XR system (Bio-Rad, USA). Densitometric analysis of the blots was performed using Fiji. Band intensities were quantified by measuring the area of each peak, with background correction applied to all measurements. Target protein levels were normalized to the corresponding β-actin loading control. Quantification was based on three independent experiments.

### Sample Preparation for Untargeted Lipidomics

MDA-MB-231 cells were seeded into 6-well culture plates at a density of 3 × 10^5^ cells per well and allowed to adhere for 24 h. Cells were then pretreated for 2 h with or without 20 μM DGAT1 and 20 μM DGAT2 inhibitors (DGATi), followed by 25 μM docosahexaenoic acid (DHA) supplementation with or without DGATi for 1, 4 and 24 h. After incubation, cells were washed twice with DPBS and harvested by scraping in 550 µl of LC/MS grade 55:45 methanol:water. Samples were stored at –80 °C until analysis. Protein content was determined in each sample using the Pierce 660 nm Protein Assay Kit following the manufacturer’s instructions and used to normalize lipid signals. The experiment included three independent biological replicates. LC/MS-grade methanol (8402-2500) was from Avantor and LC/MS-grade water (39253-1L) was from Honeywell Chemicals.

### Lipid Extraction and Untargeted LC/MS-MS Lipidomics

To extract complex lipids, we processed the samples using the LIMeX LC–MS workflow (LIpids, Metabolites, and eXposome compounds). Metabolites were extracted with a biphasic solvent system consisting of cold methanol, methyl tert-butyl ether, and 10% methanol [71]. An aliquot of the upper organic phase was collected, evaporated, and resuspended in methanol containing the internal standard [12-[(cyclohexylamino)carbonyl]amino]-dodecanoic acid (CUDA). The samples were then analyzed using lipidomics platforms in both positive and negative ion modes. For untargeted lipidomics, the LC–MS systems consisted of a Vanquish UHPLC System (Thermo Fisher Scientific, Bremen, Germany) coupled to a Q Exactive Plus mass spectrometer (Thermo Fisher Scientific, Bremen, Germany). We separated the lipids on an ACQUITY Premier BEH C18 column (50 × 2.1 mm; 1.7 μm) with VanGuard FIT (5 × 2.1 mm; 1.7 μm) and detected them using positive and negative electrospray ionization (ESI). Raw data were processed using MS-DIAL 4.94 [72] with an in-house retention time–m/z library and MS/MS libraries from public and commercial sources (MassBank, MoNA, NIST23). Data were filtered using blank samples, serial dilution samples, and quality control (QC) pool samples with a relative standard deviation (RSD) < 30%. Normalization was performed using the LOESS approach, based on QC pool samples injected regularly between every 10 actual samples. Samples were randomized across the platform run. Metabolites were annotated according to the LIPID MAPS classification system and the shorthand notation for lipid structures [73]. All analytes in MS-DIAL were manually curated.

### Lipidomic Data Analysis

Data analysis was performed using protein-normalized signal intensities (arb. units/µg protein). Outliers within biological triplicates were identified by calculating the coefficient of variation (CV); values with a CV greater than 30% were considered outliers and removed. Missing values were imputed by averaging the remaining two biological replicates. Lipids in individual samples that still exhibited a CV greater than 30% after this step were excluded from further analysis. For heatmap analysis, lipids were sorted by increasing carbon number and degree of unsaturation (number of double bonds). Lipid signal intensities were normalized and plotted on a heatmap without data clustering. For volcano plot analysis, lipid signal intensities were log_2_-transformed, and multiple unpaired *t*-tests (*P* < 0.05) were performed. To analyze relative lipid class abundance, individual lipids were grouped according to their respective lipid classes, and the summed intensities of all lipids within each lipid class were calculated for each sample. To determine the relative abundances of saturated (SFA), monounsaturated (MUFA), and polyunsaturated (PUFA)-containing triacylglycerols (TGs) and phospholipids (PLs), lipids were grouped based on their acyl-chain unsaturation. The relative abundances of all lipids within each group were summed to obtain a cumulative value, which was used to calculate the proportion of each unsaturation group relative to all TGs and PLs. The unsaturation index was calculated for each lipid by weighing the total number of double bonds by the relative abundance of that lipid within its lipid class. The individual indexes were summed for each replicate to generate a class-level unsaturation index. For lipid over-representation analysis (LORA), query lists were generated based on multiple unpaired *t*-tests (*P* < 0.1) on log_2_-transformed lipidomic data, selecting significantly upregulated lipids. Enrichment analysis was performed using the LORA online application [60]. Principal component analysis (PCA) was conducted using Metaboanalyst 5.0 to assess the distribution and consistency of the lipidomic data. GraphPad Prism 10.3.1 (Version 10, GraphPad Software Inc., USA) was used for statistical analysis, data transformation and plotting of the data.

### Statistical Analysis

All data were plotted and statistical analyses performed using GraphPad Prism 10.3.1 (Version 10, GraphPad Software Inc., USA). Data are reported as mean ± SEM, except for colocalization data from microscopy images, which are reported as median ± 95% CI. For each experiment, the number of replicates is indicated in the corresponding figure legends. Statistical significance was determined using two-tailed unpaired *t*-tests or two-way ANOVA, followed by Tukey’s multiple comparison tests. Unless otherwise stated, *P* < 0.05 was considered statistically significant.

## ACKNOWLEDGMENTS

We are grateful to Sara Jereb and Jernej Šribar for their technical help. This work was supported by the Slovenian Research Agency young researcher grant to L.P., the P1-0207 Programme and J7-1818 Project grant to T.P.; a grant from the Czech Ministry of Education, Youth and Sports [LUAUS24040] to O.K.. The authors would like to acknowledge the Metabolomics Core Facility at the Institute of Physiology of the Czech Academy of Sciences for lipidomics profiling. This publication is based upon work from COST Action EpiLipidNET, CA19105, supported by COST (European Cooperation in Science and Technology). Work in the Fedorova lab is supported by ‘‘Sonderzuweisung zur Unterstützung profilbestimmender Struktureinheiten’’ by the SMWK to TUD, TG70 by Sächsische Aufbaubank and SMWK, the measure is co-financed with tax funds on the basis of the budget passed by the Saxon state parliament (to M.F.), Deutsche Forschungsgemeinschaft (FE 1236/5-1, FE 1236/8-1 to M.F.), and Bundesministerium für Bildung und Forschung (031L0315A, DEEP_HCC and 01EJ2205A, FERROPath to M.F.).

## AUTHOR CONTRIBUTIONS

A.K. and L.P. performed most experiments, analysed data, prepared figures and wrote manuscript draft; Š.K., E.J.J. prepared samples for mass spectrometry analysis, assisted with flow cytometry and microscopy, and analysed data; N.F. performed some of the flow cytometry experiments; C.P.K. performed data analysis; K.B. and O.K. performed mass spectrometry analyses, lipidomic data analysis, assisted with experimental design, contributed ideas and revised the manuscript; M.F. and M.W. contributed ideas, lipidomics expertise and revised the manuscript. T.P. conceptualised the study, analysed data, prepared figures, and revised the manuscript.

## SUPPLEMENTARY DATA FIGURE LEGENDS

**Supplementary Figure 1.**
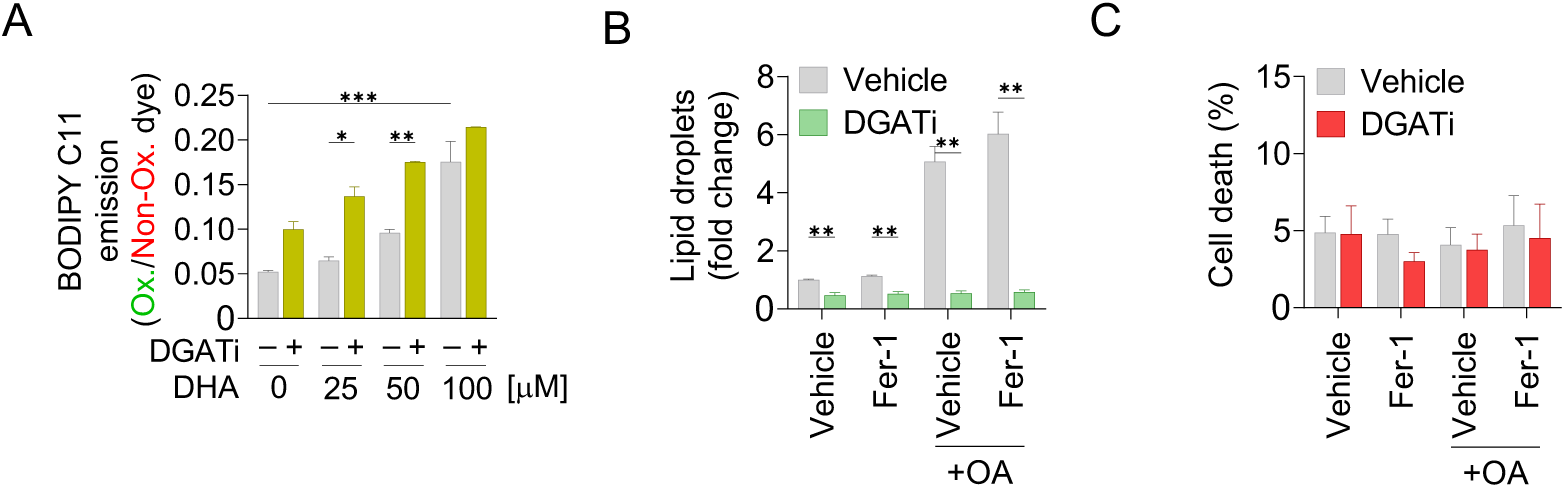
Oleic acid induces DGAT-mediated LD accumulation but not ferroptosis. (A) Lipid peroxidation assessed by BODIPY C11 581/591 probe using flow cytometry in MDA-MB-231 cells treated with 25–100 μM docosahexaenoic acid (DHA) for 24 h, with or without 20 μM DGAT1 and 20 μM DGAT2 inhibitors (DGATi). (B) Lipid droplet levels measured by Nile Red staining in MDA-MB-231 cells treated with 100 μM oleic acid (OA) and/or 5 μM ferrostatin (Fer-1) for 24 h, with or without DGATi. (C) Cell death (%) determined by TMRM/YO-PRO-1 staining in MDA-MB-231 cells following the treatment described in (B). All data are presented as mean ± SEM of at least three independent biological replications (significance calculated by two-way ANOVA with Tukey adjustment). p > 0.05 = ns; ^∗^p < 0.05; ^∗∗^p < 0.01; ^∗∗∗^p < 0.001.

**Supplementary Figure 2.**
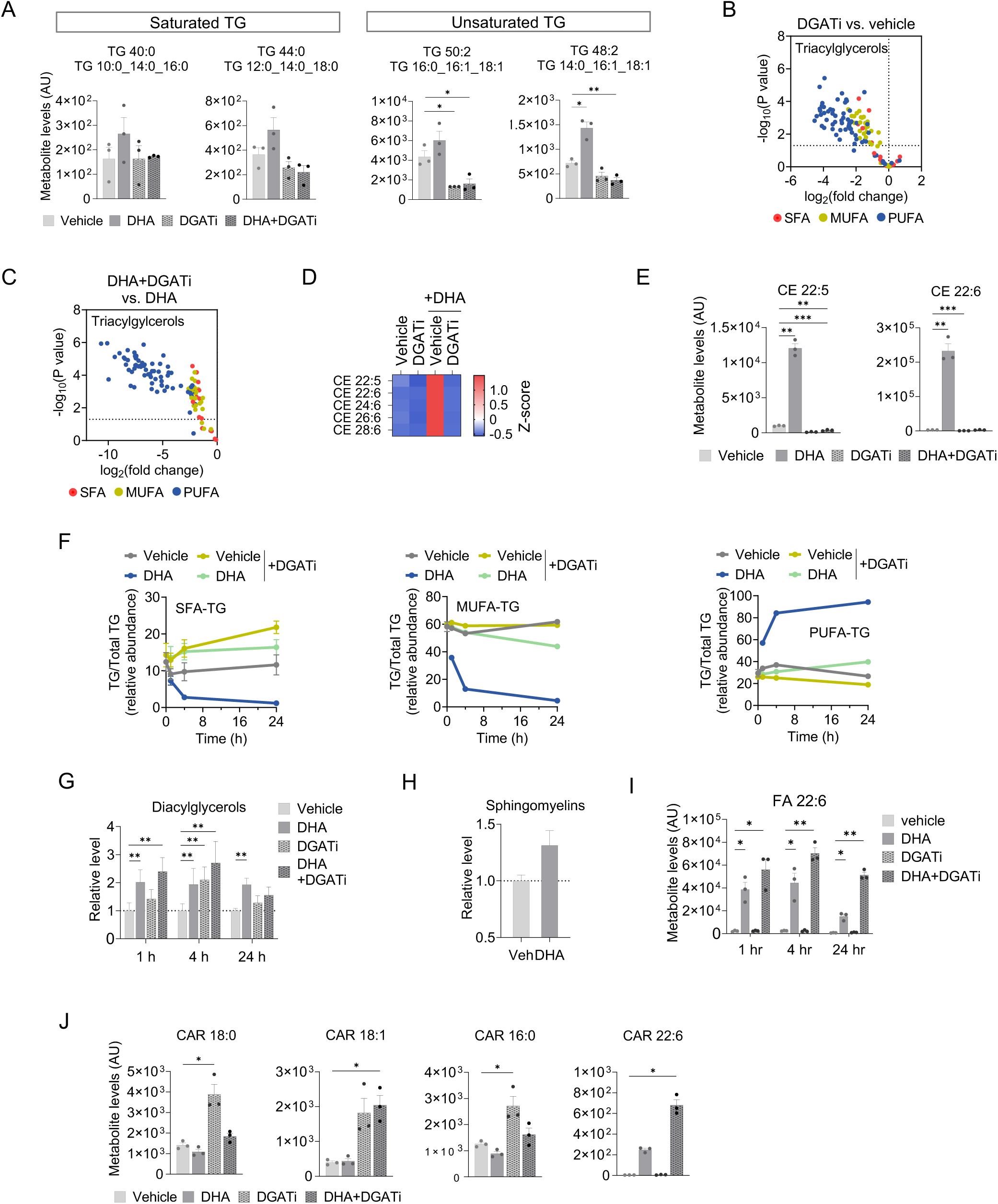
DHA and DGATi-induced changes in LD-stored neutral lipids. (A-J) MDA-MB-231 cells were treated with 25 μM docosahexaenoic acid (DHA) for 1, 4 and 24 h with or without 20 μM DGAT1 and 20 μM DGAT2 inhibitors (DGATi). At each timepoint, cellular lipids were extracted and lipidomics analysis performed. (A) Abundance of selected saturated and unsaturated TG lipids after 24 h of treatment. Each point represents a biological replicate. Data are presented as mean ± SEM (unpaired *t*-test). p > 0.05 = ns; ^∗^p < 0.05; ^∗∗^p < 0.01; ^∗∗∗^p < 0.001. (B, C) Volcano plots of TG species altered by 24 h treatment: (B) with DGATi alone, and (C) with DGATi in DHA-supplemented cells. Lipids are categorized and coloured by unsaturation as indicated in the legend. Data were generated using log_2_-transformed fold-change values and multiple unpaired *t*-tests (p < 0.05). (D) Heatmap showing relative abundance of all quantified CEs (n = 5) after 24 h of treatment. Data were Z-score normalized. (E) Selected CE lipids after 24 h of treatment. Each point represents a biological replicate. Data are presented as mean ± SEM (unpaired *t*-test). p > 0.05 = ns; ^∗^p < 0.05; ^∗∗^p < 0.01; ^∗∗∗^p < 0.001. (F) Relative abundance of SFA-, MUFA-, or PUFA-containing TGs. Data are presented as mean ± SEM of three independent biological replicates. (G) Relative total levels of diacylglycerols. Data are presented as mean ± SEM of three independent biological replicates (unpaired *t*-test). p > 0.05 = ns; ^∗^p < 0.05; ^∗∗^p < 0.01; ^∗∗∗^p < 0.001. (H) Relative total levels of sphingomyelins after 24 h of treatment. Data are presented as mean ± SEM of three independent biological replicates (unpaired *t*-test). p > 0.05 = ns; ^∗^p < 0.05; ^∗∗^p < 0.01; ^∗∗∗^p < 0.001. (I) Abundance of free DHA (FA 22:6). Each point represents a biological replicate. Data are presented as mean ± SEM (unpaired *t*-test). p > 0.05 = ns; ^∗^p < 0.05; ^∗∗^p < 0.01; ^∗∗∗^p < 0.001. (J) Abundance of selected carnitine (CAR) species after 24 h of treatment. Each point represents a biological replicate. Data are presented as mean ± SEM (unpaired *t*-test). p > 0.05 = ns; ^∗^p < 0.05; ^∗∗^p < 0.01; ^∗∗∗^p < 0.001.

**Supplementary Figure 3.**
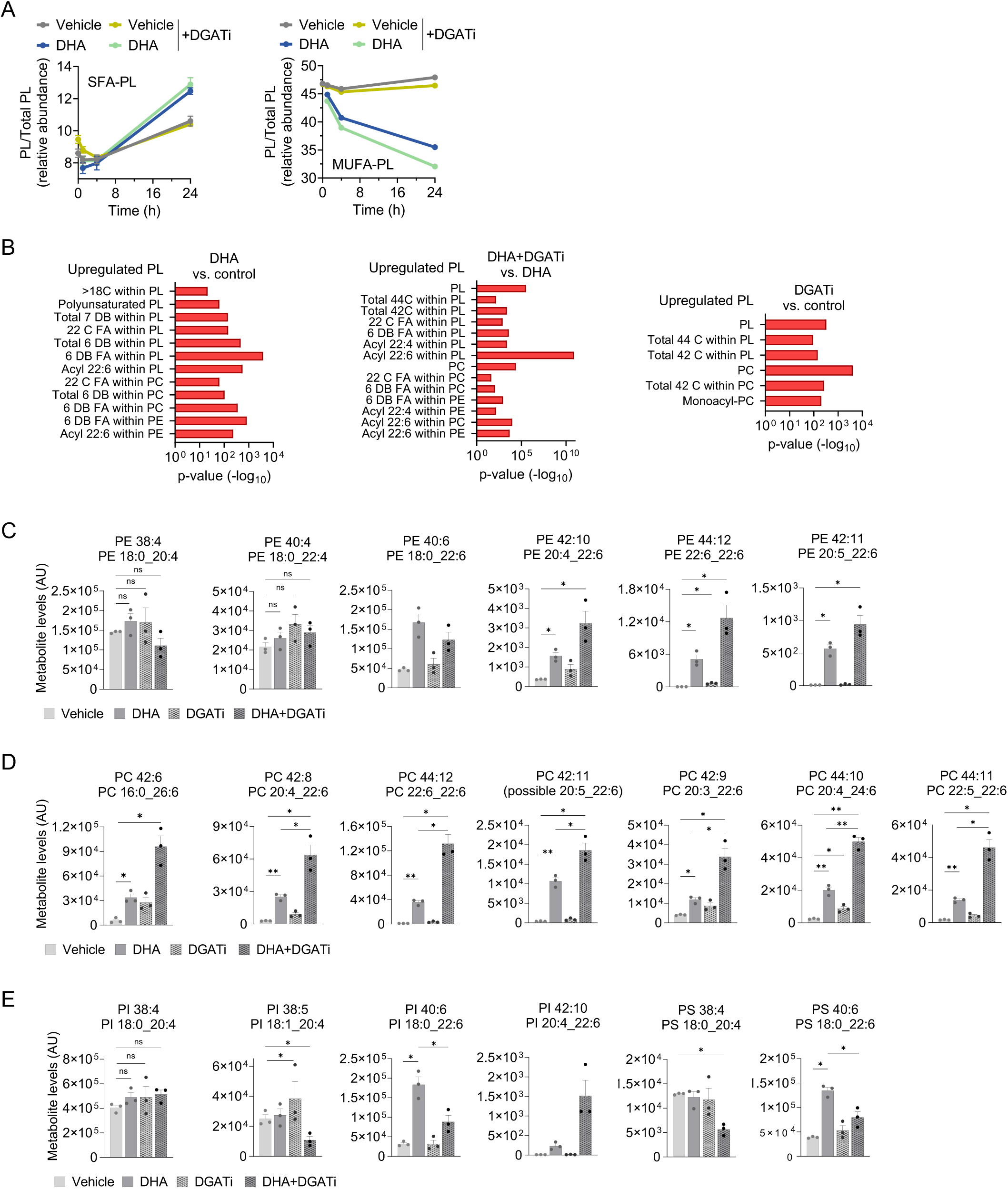
DHA and DGATi-induced changes in PL acyl-chain composition. (A) Relative abundance of SFA- and MUFA-containing PLs in MDA-MB-231 cells following 1, 4 and 24 h treatment with 25 μM docosahexaenoic acid (DHA) with or without 20 μM DGAT1 and 20 μM DGAT2 inhibitors (DGATi). Data are presented as mean ± SEM of three independent biological replications. (B) Lipid Over-Representation Analysis (LORA) of upregulated PL clusters in MDA-MB-231 cells following the 24 h treatment described in (A). (C–E) Abundance of selected PUFA-containing phosphatidylethanolamine (PE) (C), phosphatidylcholine (PC) (D), phosphatidylinositol (PI) and phosphatidylserine (PS) (E) PL species in MDA-MB-231 cells following the 24 h treatment described in (A). Each point represents a biological replicate. Data are presented as mean ± SEM (unpaired *t*-test). p > 0.05 = ns; ^∗^p < 0.05; ^∗∗^p < 0.01; ^∗∗∗^p < 0.001.

**Supplementary Figure 4.**
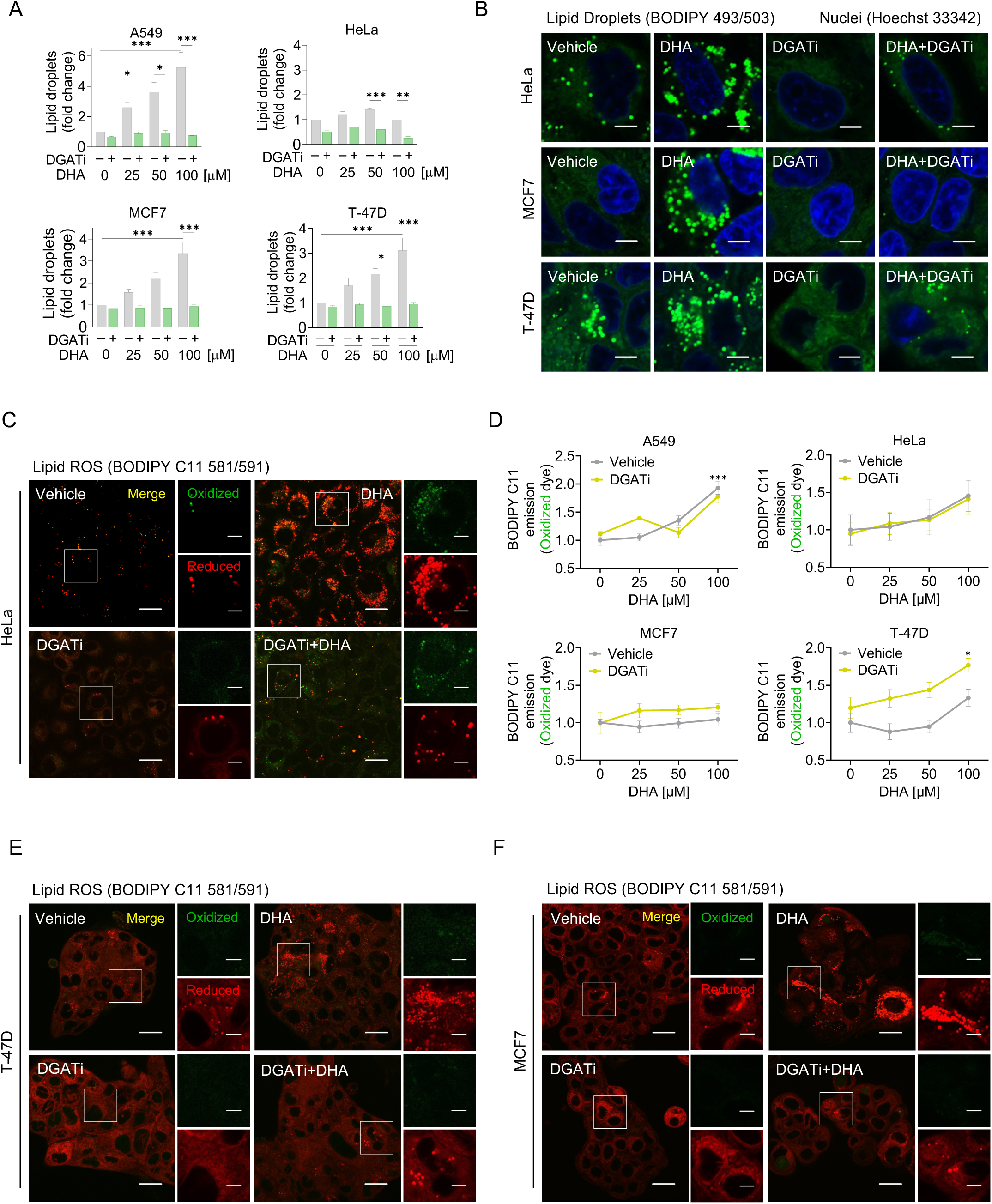
DHA- and DGATi-induced changes in LD and lipid ROS levels in ferroptosis-resistant cells. (A) Lipid droplet levels measured by Nile Red staining in indicated cell lines treated with 25-100 μM docosahexaenoic acid (DHA) for 24 h, with or without 20 μM DGAT1 and 20 μM DGAT2 inhibitors (DGATi). (B) Representative confocal microscopy images of HeLa, MCF7 and T-47D cells treated with 100 μM DHA for 24 h, with or without DGATi. Lipid droplets were stained by BODIPY 493/503 and nuclei by Hoechst 33342. Scale bar represents 5 μm. (C) Representative confocal microscopy images of HeLa cells showing BODIPY C11 581/591 lipid peroxidation induced by the treatment described in (B). Scale bar represents 20 μm and 5 μm (inset). (D) Lipid peroxidation assessed by the oxidized BODIPY C11 581/591 probe using flow cytometry in indicated cell lines following the treatment described in (A). (E, F) Representative confocal microscopy images of T-47D (E) and MCF7 (F) cells showing BODIPY C11 581/591 lipid peroxidation induced by the treatment described in (B). Scale bar represents 20 μm and 5 μm (inset). All data are presented as mean ± SEM of at least three independent biological replications (significance calculated by two-way ANOVA with Tukey adjustment). p > 0.05 = ns; ^∗^p < 0.05; ^∗∗^p < 0.01; ^∗∗∗^p < 0.001.

**Supplementary Figure 5.**
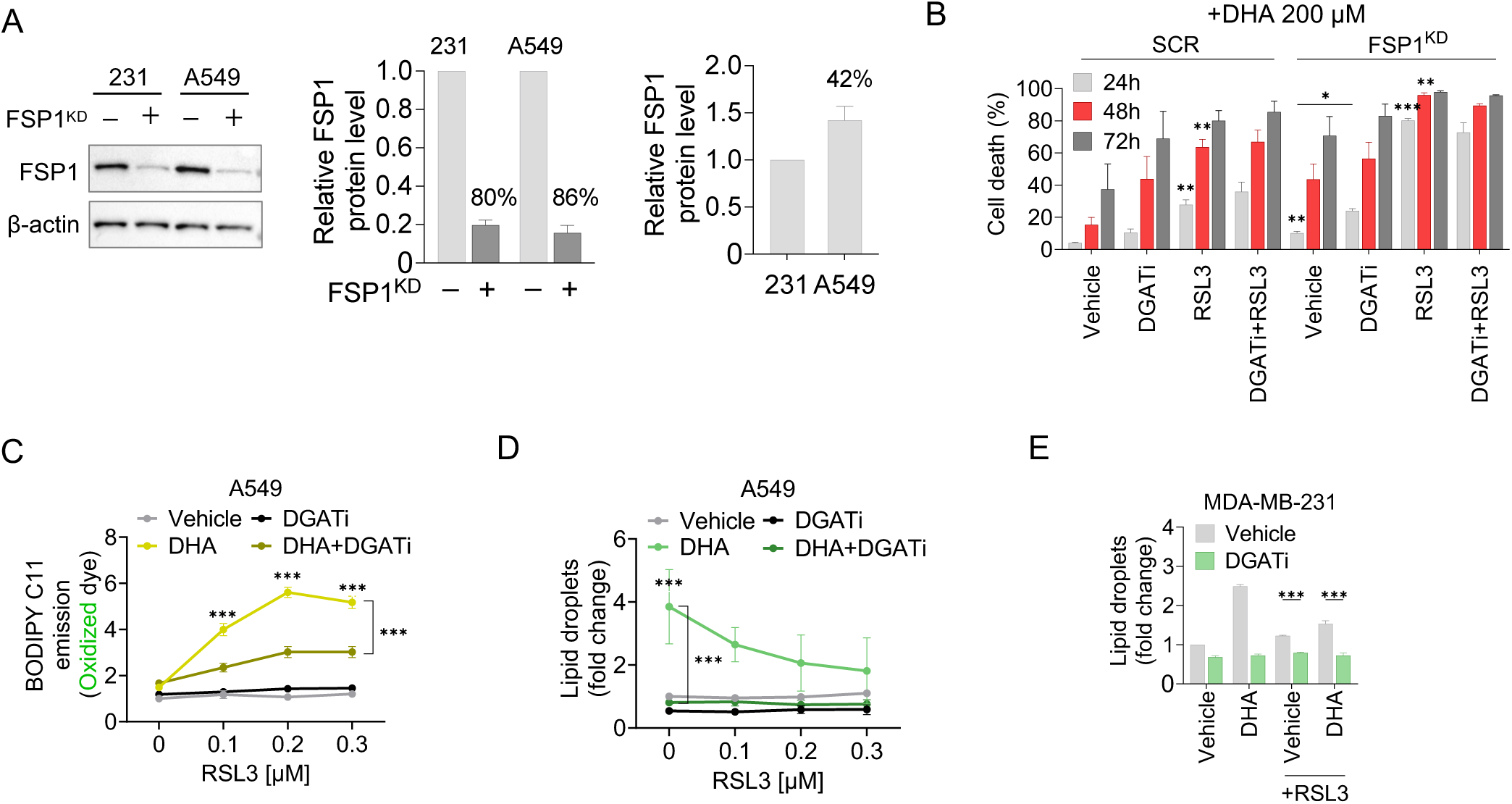
DGAT inhibition protects or promotes ferroptosis depending on the context. (A) Immunoblot detection of ferroptosis suppressor protein 1 (FSP1) and β-actin in whole-cell lysates of control (SCR) and FSP1-silenced (FSP1^KD^) MDA-MB-231 and A549 cells prepared 48 h after reverse transfection. Representative western blot is shown and densitometric analysis of n = 3 biological replicates. (B) Cell death (%) determined by TMRM/YO-PRO-1 staining in SCR and FSP1^KD^ A549 cells treated with 200 μM docosahexaenoic acid (DHA) for 24, 48 and 72 h, with or without 20 μM DGAT1 and 20 μM DGAT2 inhibitors (DGATi) and 0.1 μM RSL3. (C) Lipid peroxidation assessed by BODIPY C11 581/591 using flow cytometry in A549 cells treated with 100 μM DHA for 24 h, with or without DGATi and 0.1, 0.2 or 0.3 μM RSL3. (D) Lipid droplet levels measured by Nile Red staining in A549 cells following the treatment described in (C). (E) Lipid droplet levels measured by Nile Red staining in MDA-MB-231 cells treated with 25 μM DHA for 24 h, with or without DGATi and 0.05 μM RSL3. All data are presented as mean ± SEM of at least three independent biological replications. Significance calculated by unpaired *t*-test (B) or two-way ANOVA with Tukey adjustment (C-E). p > 0.05 = ns; ^∗^p < 0.05; ^∗∗^p < 0.01; ^∗∗∗^p < 0.001.

